# Gut Microbe-Derived *N*-Acyl Serinol Lipids Shape Host Postprandial Metabolic Homeostasis

**DOI:** 10.1101/2025.04.11.648385

**Authors:** Sumita Dutta, Kala K. Mahen, William J. Massey, Venkateshwari Varadharajan, Amy C. Burrows, Anthony J. Horak, Marko Mrdjen, Nour Mouannes, Danny Orabi, Lucas J. Osborn, Treg Grubb, Rachel Hohe, Rakhee Banerjee, Vinayak Uppin, Dev Laungani, Grace Hamilton, Xiayan Ye, Naseer Sangwan, Mohammed Dwidar, Adeline M. Hajjar, Belinda Willard, Michael Martin, Erik Guetschow, Patrick Westcott, Zeneng Wang, Stanley L. Hazen, J. Mark Brown

**Author notes:** Authors contributed equally as first author on this body of work (Sumita Dutta, Kala K. Mahen, William J. Massey). To whom correspondence should be addressed: Department of Cancer Biology, Cleveland Clinic, Cleveland, OH 44195, USA.

## Abstract

Although strong evidence links the gut microbiome to metabolic disease, the mechanisms linking microbiota to hormonal and metabolic responses to food are not well understood. After a meal, gut bacteria produce a wide array of small molecule, protein, and lipid metabolites originating from bacterial sources. Annotating physiological function to select gut microbe-derived metabolites is critical to understanding diet-microbe-host interactions, and to developing microbiome-inspired therapies to improve human health. Here, we have investigated the role of a poorly annotated class of gut microbiome-derived lipids called *N*-acyl amides in postprandial metabolic physiology. Here we show both bacterial overproduction and provision of exogenous *N*-acyl amides reorganize host hormone-driven metabolic transition after a meal. Moreover, *N*-acyl amides exert broad effects on the meal- and circadian-related reorganization of gene expression, metabolic hormones, and gut microbiome composition. Collectively, these results demonstrate that microbiota-derived *N*-acyl amides play a physiologic role in postprandial metabolic homeostasis in the host.

## INTRODUCTION

Common endocrine and metabolic disorders such as obesity, diabetes, thyroid abnormalities, polycystic ovary disease (PCOS), and osteoporosis are driven by either defective hormone production or hormone action in target tissues. Although the field of endocrinology has historically focused on understanding host organ systems responsible for hormone production (hypothalamus, pituitary, pineal, thyroid, pancreas, adipose, etc.), the gut microbiome also fits the definition of an endocrine organ with unparalleled hormone signaling potential^1–4^. Unlike host endocrine glands, which produce only a few key hormones, the gut microbial endocrine organ has the unique potential to produce hundreds if not thousands of hormone-like products including proteins/peptides, small molecules, sugars/oligosaccharides, and lipids/glycolipids^1–4^. Much like hormones derived from host endocrine organs, bacterially-derived signals are sensed by highly selective host receptor systems that elicit diverse biological responses linked to host metabolism.^1–4^ The current list of bacterially-derived metabolites that can exert endocrine properties in the host includes but is not limited to short chain fatty acids (SCFA)^5,6^, trimethylamine-N-oxide (TMAO) and related metabolites^7–9^, hydrogen sulfide^10,11^, secondary bile acids^12,13^, oligosaccharides^14^, polyamines^15^, isoflavones^16^, lignans^17^, both indole and phenolic compounds^18–24^, and lipids^25–26^. Although we are rapidly gaining an understanding of the chemical diversity originating from the gut microbial endocrine organ, there are only a handful of discoveries in the literature that mechanistically link gut microbial metabolite production to host hormonal control of metabolism.

It is important to remember that our microbiome is dynamically altered by the food we consume^27–29^. In parallel, many metabolic hormones are also released from host endocrine organs when we eat^28–30^. Given the close interactions between our dietary intake, gut microbiota, and host hormone release, it is possible that endocrine circuits exist whereby signals arising from bacteria can impact host hormone release or action. In fact, advancing understanding of diet-microbe-host endocrine circuits would have tremendous untapped therapeutic potential to combine therapies targeted to balance both microbe- and host-derived hormones.

Undoubtedly, the most exciting discovery in host endocrinology and metabolism in the past decade is the discovery that GLP-1 receptor agonist drugs can promote profound weight loss and protection from many obesity-related disorders such as type 2 diabetes, cardiovascular disease (CVD), kidney disease, and several other comorbid conditions^33–36^. GLP-1 is a 30 amino acid incretin peptide that is predominantly secreted by the distal ileal and colonic *L*-cells following a meal, and can exert diverse metabolic effects by enhancing insulin secretion, reducing food intake and delayed gastric emptying, and dampening stress-related and inflammatory pathways^33–36^. Although GLP-1 is solely host derived, there is strong evidence that the gut microbiome can powerfully impact both the production and turnover of this critical metabolic hormone^37–40^. For example, mice raised under germ-free conditions or treated with broad spectrum antibiotics have elevated levels of active GLP-1^37,38^. Furthermore, gut bacteria encode a GLP-1-cleaving dipeptidyl peptidase that can degrade host-derived GLP-1 to shape meal-related incretin responses^39^. Also, an independent report showed that a novel class of bacterially-derived lipids called *N-*acyl amides can stimulate production of GLP-1 by activating host GPR119 on enteroendocrine *L*-cells^40^. *N-*acyl amides are produced by gut microbial *N-*acyl synthase (NAS) enzymes, and the *nas* genes encoding this enzymatic capacity have been identified in human fecal samples^40^. In initial studies colonizing germ-free mice with gut bacteria encoding *nas* genes, it was demonstrated that bacterially-derived *N*-acyl amides can improve glucose tolerance^40^. Although this study showed initial support that bacterial *N*-acyl amides can improve host hormonal control of metabolism, many questions remained unanswered. Here, we explore how elevation of *N*-acyl amide lipid levels impact postprandial hormones and metabolic homeostasis. Our studies demonstrate that gut microbe-derived *N*-acyl serinol lipids exert broad effects on the meal- and circadian-related reorganization of gene expression, metabolic hormones, and gut microbiome composition in mice.

## RESULTS

### *N*-Oleoyl Serinol Reorganizes Host Hormonal and Metabolic Responses Following a Meal

To study the effects of *N*-acyl serinol lipids on meal-related physiology, we initially synthesized sufficient quantities of *N*-oleoyl serinol and incorporated it into slow-release pellets that allow for long-term systemic release *in vivo*^18,41^. In parallel, we also generated a stable isotope dilution liquid chromatography tandem mass spectrometry (LC-MS/MS) method to quantitate circulating and tissue levels of microbe- and host-derived *N*-acyl amide lipids. In the first set of studies we implanted slow-release pellets to elevate *N-*acyl amides and then collected tissues under ad libitum fed, fasted, or fasted and then re-fed conditions to understand their effects on hormonal control of metabolism (Figures 1, S1, and S2). As intended, slow-release pellets raised plasma *N*-oleoyl serinol ∼2-3 fold (Figure 1A) but did not significantly alter circulating levels of other microbe- or host-related fatty acid amide lipids such as *N*-palmitoyl serinol (Figure 1B), palmitoylethanolamide (PEA) (Figure 1C), stearoylethanolamide (SEA) (Figure 1D), or oleoylethanolamide (OEA) (Figure 1E). It is important to note that in control mice plasma *N*-oleoyl serinol levels did not change with fasting or feeding, but in the liver of control mice fasting significantly reduced levels and re-feeding promoted a 3-fold increase in *N*-oleoyl serinol levels indicating meal responsiveness (Figure S1A).

**Figure 1.**
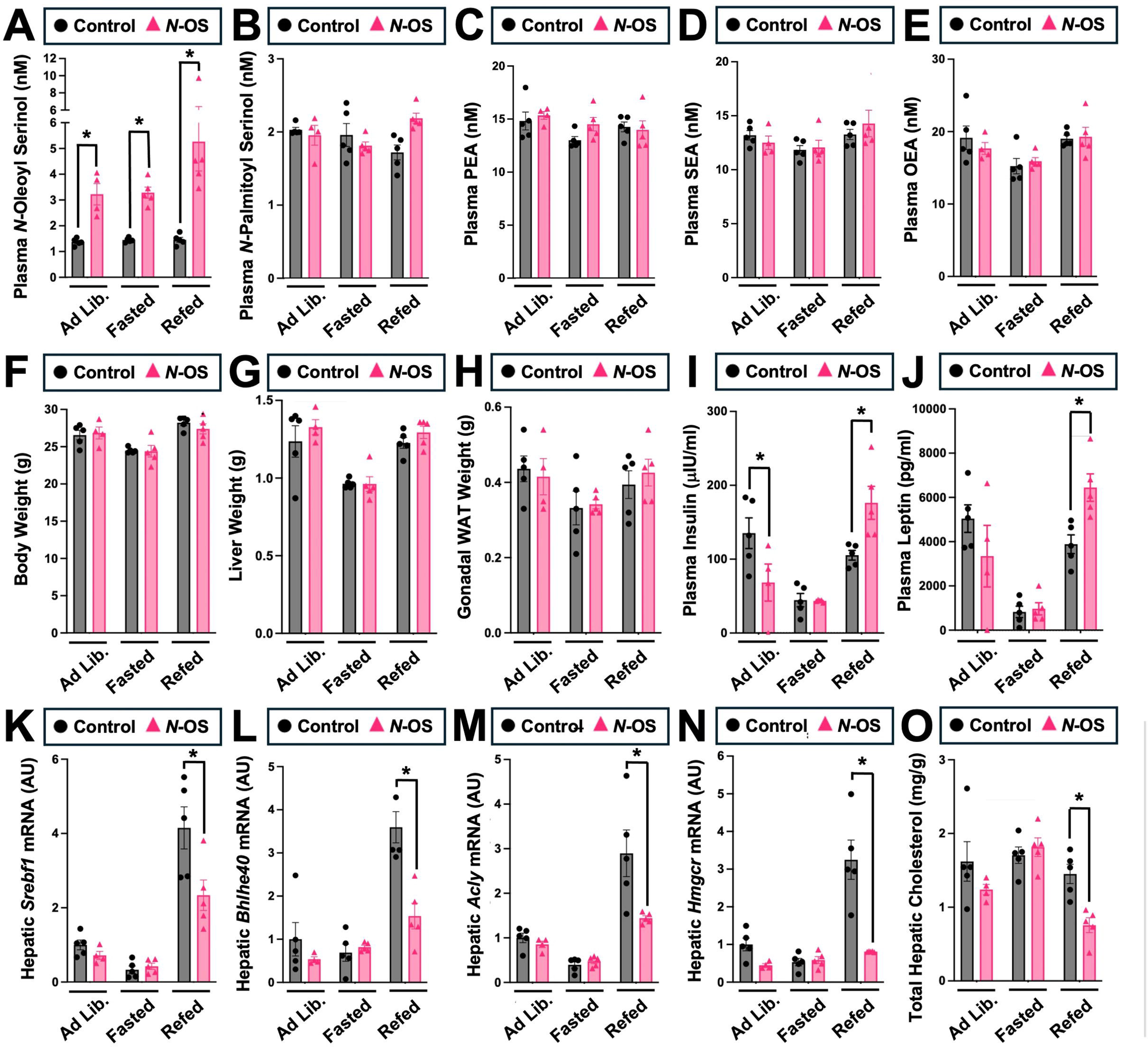
*N*-Oleoyl Serinol Alters Postprandial Metabolic Homeostasis. Male chow-fed C57Bl/6J mice were implanted with slow-release pellets containing empty scaffold (control) or *N*-oleoyl serinol (*N*-OS). One week later, mice were necropsied under ad libitum feeding (Ad Lib.), 12 hours fasted (Fasted), or 12 hours fasted and then re-fed for 3 hours (Re-Fed) to examine meal-related hormone alterations. (**A-E**) Plasma levels of microbe and host derived acylamide lipids including *N*-oleoyl serinol (*N*-OS), *N*-palmitoyl serinol (*N*-PS), palmitoylethanolamide (PEA), oleoylethanolamide (OEA), and arachidonoylethanolamide (AEA) were quantified by LC-MS/MS. Body weight (**F**), liver weight (**G**), and gonadal white adipose tissue (gWAT) weight (**H**) were measured at necropsy. (**I,J**) Plasma levels of insulin and leptin were quantified by immunoassay. (**K-O**) The relative gene expression for metabolism related genes in the liver was quantified by qPCR. Data represent the mean ± S.E.M. from n=4-5 per group, and statistically significant difference (Student’s t-tests) within each feeding status group are denoted by * = p<0.05.

*N*-oleoyl serinol treatment did not significantly alter body weight (Figure 1F), liver weight (Figure 1G), or gonadal adipose tissue weight (gWAT) (Figure 1H), but did significantly alter meal-related hormonal and metabolic responses systemically. Mice receiving *N*-oleoyl serinol pellets have unaltered fasting/re-feeding responses in hepatic (Figure S1B) and plasma (Figure S1D) triglyceride levels, but did have modestly elevated plasma total cholesterol and phospholipid levels in the ad libitum-fed state (Figure S1E, S1F). *N*-oleoyl serinol treatment also elevated hepatic phospholipid levels only under fasted conditions (Figure S1C). Unexpectedly, we found GLP-1 levels were reduced in mice receiving *N*-oleoyl serinol in the ad libitum-fed state (Figure S1G), and alterations in several other meal-related hormones were altered in mice receiving exogenous *N*-oleoyl serinol (Figures 1I, 1J, S1G-I). *N*-oleoyl serinol treatment reduced plasma insulin in the ad libitum fed state, but reciprocally increased insulin levels in the refed state (Figure 1I). Mice receiving *N*-oleoyl serinol also had significantly elevated plasma leptin levels in the refed state (Figure 1J). Only in the ad libitum fed state, mice treated with *N*-oleoyl serinol had reduced levels of peptide YY (PYY) and glucagon (Figure S1H, S1I). However, *N*-oleoyl serinol did not significantly alter plasma cytokine levels including interleukin 6 (IL-6), monocyte chemoattractant protein 1 (MCP-1) and tumor necrosis factor α (TNFα) (Figure S1J-S1L). Collectively, these results demonstrate that *N*-oleoyl serinol can have diverse effects on select meal-related hormone levels.

Postprandial hyperinsulinemia and hyperleptinemia are commonly associated with obesity and prediabetes, but this is most often coupled with insulin and leptin resistance in target tissues^42,43^. Despite elevations in postprandial insulin and leptin observed with *N*-oleoyl serinol treatment (Figure 1I, IJ), the postprandial insulin-driven response in lipogenic gene expression was blunted. For instance, several insulin-responsive genes involved in *de novo* lipogenesis, including sterol regulatory element-binding protein 1 and 2 (SREBP1 and SREBP2) targets, were reduced in mice receiving *N*-oleoyl serinol in the re-fed state where insulin is most active (Figure 1K-1N). In the re-fed state, *N*-oleoyl serinol treatment was associated with reduced expression of insulin-driven lipogenic genes, including basic helix-loop-helix family member E40 (*Bhlhe40*)^44^ (Figure 1L), and ATP-citrate lyase (*Acly*) (Figure 1M). Furthermore, *N*-oleoyl serinol treatment markedly suppressed the expression of HMG-CoA reductase (*Hmgcr*), which was associated with reduction in hepatic cholesterol in the refed state (Figure 1O). It is important to note that *N*-oleoyl serinol treatment did not alter the hepatic expression of glucose-related genes including phosphoenolpyruvate carboxykinase (*Pck1*) and glucose 6 phosphate dehydrogenase (*G6pd*) (Figure S1M, S1N). Furthermore, expression of fibroblast growth factor 21 (*Fgf21*) and carnitine palmitoyltransferase 1α (*Cpt1*α) were unaffected by *N*-oleoyl serinol treatment (Figure S1O). However, *Cpt1*α expression was reduced in the liver of mice receiving *N*-oleoyl serinol in the refed state (Figure S1P). Also, hepatic expression of the genes encoding cholesterol-esterifying enzymes for acyl-CoA:cholesterol acyltransferases 1 and 2 (*Soat1* and *Soat2*) were unaffected in the ad libitum and fasted state, but *Soat2* was reduced in the refed state in *N*-oleoyl serinol-treated mice (Figure S1Q, S1R). When we examined metabolic gene expression in gonadal white adipose tissue (gWAT), the only gene monitored that was significantly altered was *Cpt1*α, which was reduced in *N*-oleoyl serinol-treated mice only in the fasted state (Figure S1S-S1X).

In an effort to further explore the impact of *N*-oleoyl serinol effect on host metabolism we performed unbiased proteomics in livers from *N*-oleoyl serinol-treated mice. Notably, the *N*-oleoyl serinol induced proteomic alterations observed were dependent on the metabolic state of the mice. In the ad libitum condition, *N*-oleoyl serinol treatment strikingly downregulated proteins known to regulate the spliceosome, oxidative phosphorylation, and thermogenesis, yet upregulated proteins involved in AMPK signaling, bile acid synthesis, and glucagon signaling (Figure S2A-S2C, and Supplemental Table S2). Under fasted conditions, *N*-oleoyl serinol treatment downregulated proteins related to Yersinia infection, peroxisome function, and PD-L1/PD-1 checkpoint pathways, and upregulated proteins involved in ribosome function, coronavirus infection, and diabetic cardiomyopathy (Figure S2D-S2F, and Supplemental Table S2). In contrast, in the refed state, *N*-oleoyl serinol treatment suppressed proteins involved in ribosome function, coronavirus disease, and spliceosome function, and upregulated proteins involved in AMPK and glucagon signaling, and RNA transport (Figure S2G-S2I, and Supplemental Table S2).

### *N*-Oleoyl Serinol Alters Meal-Related Reorganization of the Gut Microbiome

Bi-directional microbe-host communication is required for homeostatic control of meal-related physiology in the metaorganism^45–48^. We next tested whether *N*-oleoyl serinol affected meal-related alterations in the gut microbiome itself. Principal coordinates analysis of microbial taxa revealed distinctly differentiated clusters between control and *N*-oleoyl serinol-treated cecal microbiomes under the ad libitum fed, fasted, and refed states (Figure 2A). In particular, the meal-related alterations in several genera, including *Bacteroides*, *Fecalibacterium*, *Blautia*, *Lachnospiraceae, Dubosiella*, *Turicibater*, *Alistipes*, and *Akkermansia*, are observed in *N-*oleoyl serinol-treated mice compared to controls (Figures 2B). The most significantly altered taxa under the ab libitum fed state included *Romboutsia*, *Clostridium sensu stricto 1*, *Defluviitaleaceae UCG-011*, and *Eubacterium nodatum* group (Figure 2C). In the fasted state, *Romboutsia* and *Clostridium sensu stricto* continued to be altered by *N*-oleoyl serinol treatment, but *Dubosiella* was also strikingly reduced in *N*-oleoyl serinol-treated mice (Figure 2D). In the refed state, the most differentially abundant taxa in *N*-oleoyl serinol treated mice included *Alistipes* and *Turicibacter* (Figure 2E). Although there were many cecal microbiome alterations in *N*-oleoyl serinol-treated mice, we wanted to investigate which genera were correlated with hormonal and metabolic alterations in the host. The genera that were most significantly correlated with plasma insulin levels included *ASF356*, *Akkermansia*, *Bifidobacterium*, *UCG-005*, and *Blautia* (Figure 2F). However, the genera most correlated with hepatic lipogenic gene expression (*Srebf1* and *Hmgcr*) were *Anaerotruncus*, *Bifidobacterium*, *CAG-56*, *Blautia*, and *Coprococcus* (Figure 2G, 2H). These data indicate that in addition to altering metabolic physiology in the host, systemic elevation of *N*-oleoyl serinol in the host also impacts meal-related reorganization of the gut microbiome.

**Figure 2.**
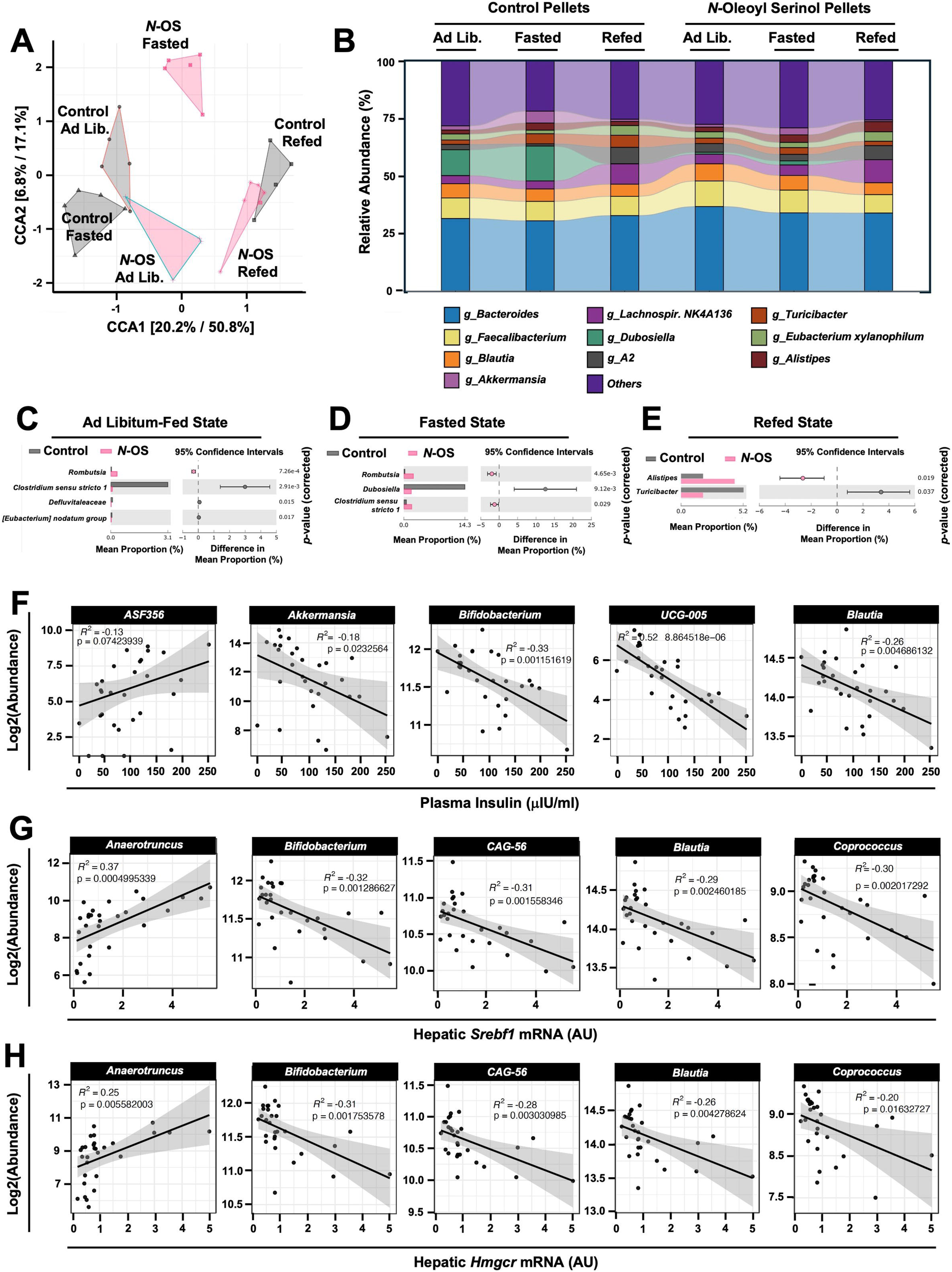
*N*-Oleoyl Serinol Shapes Meal-Related Reorganization of the Gut Microbiome. Male chow-fed C57Bl/6J mice were implanted with slow-release pellets containing empty scaffold (control) or *N*-oleoyl serinol (*N*-OS). One week later, mice were necropsied under ad libitum feeding (Ad Lib.), 12 hours fasted (Fasted), or 12 hours fasted and then re-fed for 3 hours (Re-Fed) to collect cecum for microbiome composition analyses via sequencing the V4 region of the 16S rRNA (genus level changes are shown). **(A)** Canonical correspondence analysis (CCA) based beta diversity analyses show distinct microbiome compositions in *N*-oleoyl serinol treated mice under defined fed or fasted conditions. Statistical significance and beta dispersion were estimated using PERMANOVA. **(B)** The relative abundance of cecal microbiota in control and *N*-oleoyl serinol treated mice respond differently during fasting and feeding transitions. Significantly altered cecal microbial genera in control and *N*-oleoyl serinol treated mice are shown in the ad libitum fed state (**C**), fasted state (**D**), or fasted and then refed state (**E**); ASVs significantly different in abundance (MetagenomeSeq with Benjamini-Hochberg FDR multiple test correction, adjusted P % 0.01). (**F-H**) Correlation analyses comparing the abundance of ASVs and plasma insulin (**F**), hepatic *Srebf1* mRNA expression (**G**), or hepatic *Hmgcr* mRNA expression (**H**). Data shown represent the means ± S.D. for n= 5 individual mice per group. Group differences were determined using ANOVA with Benjamini-Hochberg false discovery rate (FDR).

### Gnotobiotic Mouse Studies Further Demonstrate that Gut Microbe-Derived *N*-Acyl Amides Shape Fasting/Re-Feeding Reponses in the Host

Although slow-release pellets have clear utility, peripheral systemic administration with this approach does not recapitulate the endogenous bacterial production of *N*-acyl amides in the gut. To specifically understand the role of gut microbe-derived *N*-acyl amides in host metabolic physiology, we cloned the bacterial *nas* gene from Gemella haemolysans M34^40^ and stably expressed it in a mouse colonizing *E. coli* strain (MP1). Wild type (WT) *E. coli* does not harbor any predicted *nas* genes, and in fact does not produce any detectable *N*-acyl amides in culture, yet our *nas*-expressing *E. coli* produce ample amounts of *N*-oleoyl serinol (Figure 3A). Next, we monocolonized wild type C57BL/6N germ-free mice with either WT or NAS-expressing *E. coli* and found significantly elevated levels of multiple bacterially-derived *N*-acyl amides, including *N*-oleoyl serinol, *N*-palmitoyl serinol, and *N*-arachidonoyl serinol in gnotobiotically housed mice (Figure 3B-3D). Under the fasted and fed conditions, we did not see significant alterations in active GLP-1 in mice harboring *nas*-expressing *E. coli* (Figure 3E). Control mice harboring WT *E. coli* showed significant reductions in circulating ghrelin after fasting, but this fasting-induced suppression of ghrelin was not apparent in mice colonized with *nas*-expressing *E. coli* (Figure 3F). Furthermore, mice colonized with *nas*-expressing *E. coli* had significantly elevated plasma leptin and MCP-1 levels specifically in the fasted state (Figure 3G, 3H). Also, the expression of the fibroblast growth factor 21 (*Fgf21*) gene was dramatically increased in mice colonized with *nas*-expressing *E. coli* under fed conditions, and the lipogenic gene acetyl-CoA carboxylase (*Acaca*) was increased in mice colonized with *nas*-expressing *E. coli* only under fasted conditions (Figure 3J). We also found that fasting elicited reduced expression of fatty acid synthase (*Fasn*) in mice harboring *nas*-expressing *E. coli*, but not in mice monocolonized with WT *E. coli* (Figure 3K). The expression of ATP-citrate lyase (*Acly*) was induced by fasting in mice harboring *nas*-expressing *E. coli*, but not in mice monocolonized with WT *E. coli* (Figure 3L).

**Figure 3.**
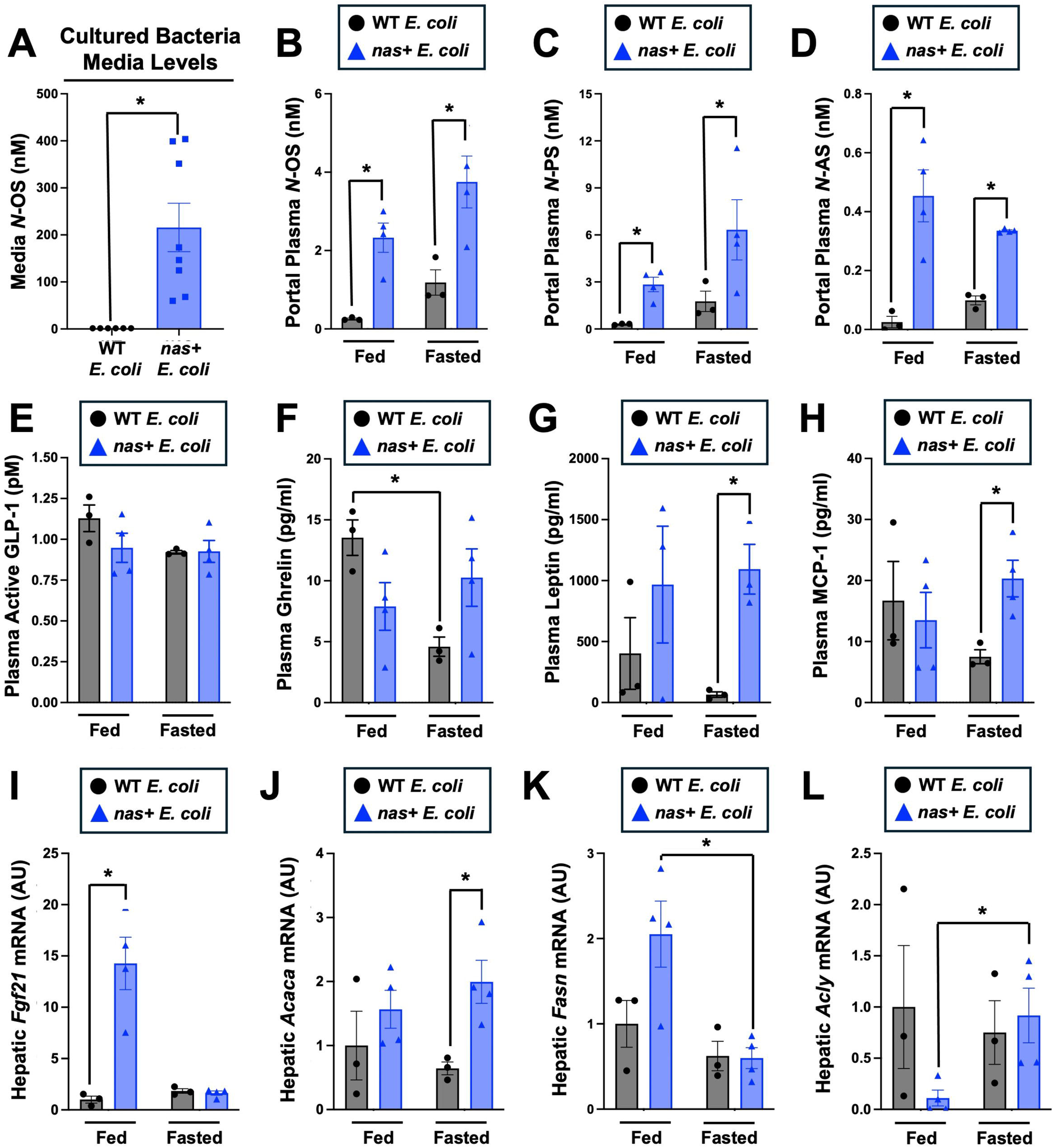
The Bacterial Gene N-Acyl Synthase (*nas*) Alter Meal-Related Metabolic Homeostasis in the Host. The *nas* gene was cloned from *Gemella haemolysans* M34^20^ and expressed in a mouse colonizing *E. coli* strain (MP1). **(A)** Wild type (WT) and *nas*-expressing (*nas+*) *E. coli* were isolated from monocolonized mice and grown overnight in LB media and media levels of *N*-oleoyl serinol were quantified by LC-MS/MS. For mouse gnotobiotic studies (B-L), germ-free C57LB/6N mice were monocolonized with either WT or *nas*-expressing *E. coli* and then 3 weeks later necropsied under the ad libitum fed state or fasted state. (**B-C**) *N*-oleoyl serinol (*N*-OS), and *N*-palmitoyl serinol (*N*-PS), and *N*-arachidonoyl serinol (*N*-AS) were quantified by LC-MS/MS using plasma collected from the portal vein. Peripheral plasma levels of GLP-1 (**E**), ghrelin (**F**), leptin (**G**), and monocyte chemoattractant protein 1 (MCP-1) were quantified by MesoScale U-Plex assay. **(I-L)** Hepatic mRNA expression for metabolic genes including (**I**) fibroblast growth factor 21 (*Fgf21*), (**J**) acetyl-CoA carboxylase α (*Acaca*), (**K**) fatty acid synthase (*Fasn*), and (**L**) ATP-citrate lyase (*Acly*) were quantified by qPCR. Data represent the mean ± S.E.M. from n=5 per group, and statistically significant difference (Student’s t-tests) are denoted by * = p<0.05.

We next performed unbiased proteomics on the liver of mice monocolonized with WT or *nas*-expressing E. coli (Figure S2J-S2O). Under fasted conditions, monocolonization with *nas*-expressing *E. coli* resulted in downregulation of proteins related to thermogenesis, glycan, fatty acid, and branched-chain amino acid (BCAA) degradation, and upregulation of proteins related to shigellosis, sulfur amino acid metabolism, and longevity (Figure S2J-S2L, and Supplemental Table S2). Under fed conditions, monocolonization with *nas*-expressing *E. coli* resulted in downregulation of proteins related to bacterial invasion of epithelial cells, actin cytoskeleton, and ribosomal biogenesis, and upregulation of proteins related to sulfur amino acid metabolism, fatty acid degradation, and thermogenesis (Figure S2M-S2O, and Supplemental Table S2).

### *N*-Oleoyl Serinol Broadly Reorganizes Circadian Rhythms in Systemic Metabolism

Given that we established that both exogenous (Figures 1, 2, S1, and S2) and bacterially-derived *N*-acyl amides (Figures 3, S2, and Supplemental Table S2) can alter hormonal and metabolic physiology under defined meal conditions (ad libitum fed, fasted, or refed), we next wanted to study whether *N*-oleoyl serinol can also shape systemic metabolic physiology over a 24-hour circadian cycle where multiple meals and core circadian clock machinery of the host intersect to shape metabolism in the metaorganism. Here, we elevated systemic *N*-oleoyl serinol levels using slow-release pellets and subsequently analyzed the effects on the gut microbiome and host hormone and metabolic homeostasis over a 24-hour (12:12 light:dark) circadian period (Figures 4, S3 and S4). As intended, slow-release pellets were able to maintain high levels of plasma *N*-oleoyl serinol throughout the 24-hour period (Figure 4A). It is also important to note that in mice receiving control pellets, there was a clear circadian rhythmic pattern of plasma *N*-oleoyl serinol that peaked in the early dark cycle where mice are most active and eating (Figure 4A). In this circadian experiment *N*-oleoyl serinol treatment resulted in a profound phase shift in plasma GLP-1 that was characterized by reduced levels in the light cycle but increased levels during the dark cycle compared to control mice (Figure 4B). Elevation of *N*-oleoyl serinol also significantly altered the circadian oscillations of plasma leptin, TNFα, and MCP-1 (Figure 4C-4E). These diverse effects on host hormone levels occurred in parallel to striking reorganization of circadian rhythms in host gene expression and metabolic homeostasis. For example, the amplitude of hepatic gene expression of key circadian clock regulators REV-ERBα (*Nr1d1*) and cryptochrome 2 (*Cry2*) was reduced in *N*-oleoyl serinol treated mice (Figure 4F, 4G). In parallel, the circadian rhythmic expression of key host metabolic genes including *Cpt1α*, *Srebf2*, and *G6pd* was suppressed in the liver of *N*-oleoyl serinol treated mice (Figure 4H-4J).

**Figure 4.**
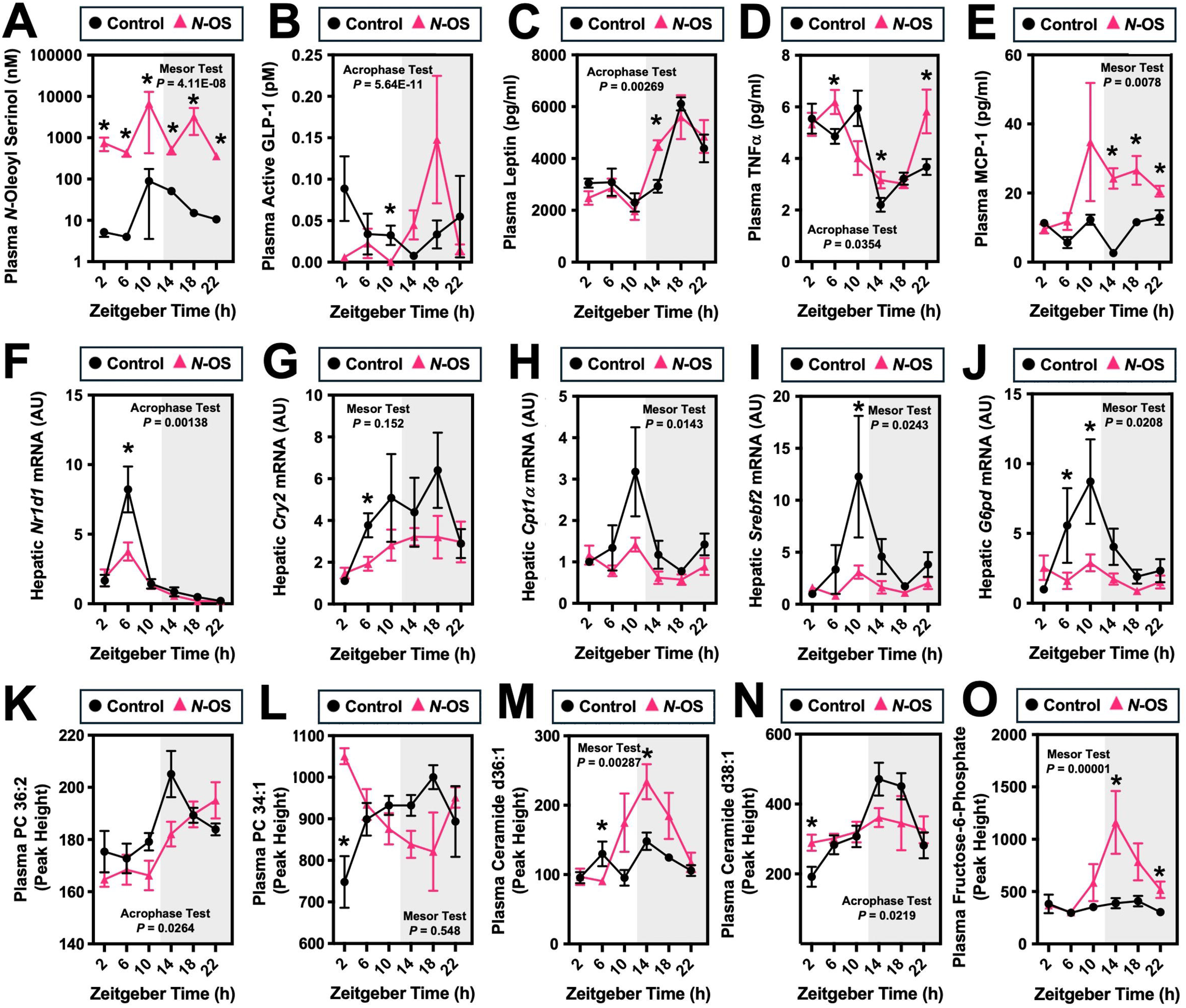
N-Oleoyl Serinol Alters Circadian Rhythms in Metabolism. Chow-fed C57Bl/6J mice were implanted with slow-release pellets containing empty scaffold or *N*-oleoyl serinol (*N*-OS). One week later, mice were necropsied over a 24-hour period (every 4 hours) to examine circadian alterations in metabolic homeostasis. (**A**) Plasma levels of N-oleoyl serinol were quantified using LC-MS/MS. (**B-E**) Plasma levels of hormones (glucagon-like peptide 1, GLP-1 and leptin) and cytokines (tumor necrosis factor α, TNFα; monocyte chemoattractant 1, MCP-1) were quantified using immunoassay. Hepatic mRNA expression for genes including (**F**) REV-ERBa (*Nr1d1*), (**G**) cryptochrome 2 (*Cry2*), (**H**) carnitine palmitoyltransferase 1 α (*Cpt1α*), (**I**) sterol regulatory element-binding transcription factor 2 (*Srebf2*), and (**J**) glucose-6-phosphate dehydrogenase (*G6pd*) were quantified by qPCR. (**K-O**) Plasma levels of metabolites showing altered circadian rhythms were quantified by either LC-MS or GC-MS. Data represent the mean ± S.E.M. from n=5 per group. Representative group differences were determined using cosinor analyses *p*-values are provided where there were statistically significant differences between control and N-oleoyl serinol treated mice. The entire analysis for circadian cosinor statistics can be found in Table S1. * = statistically significant difference (*p*<0.05) between groups within each ZT time point (Student’s t-tests).

Next, we performed untargeted metabolomics to unbiasedly identify the impact of *N*-oleoyl serinol treatment on circulating metabolite fluctuations over a 24-hour circadian period (Figures 4K-4O, and S4). The circadian oscillation in plasma phosphatidylcholine (PC) levels were altered in a molecular species-specific manner (Figures 4K, 4L, and S3A-S3F). Although the oscillations of many species of PC were unaltered by *N*-oleoyl serinol treatment (Figure S3A), the circadian oscillation of many other PC species were either enhanced or suppressed by exogenous *N*-oleoyl serinol (Figures 4K, 4L, and S3A-S3F). In parallel, *N*-oleoyl serinol treatment selectively altered molecular species of ceramides, with many unchanged and others exhibiting altered mesor or amplitude aspects of circadian rhythms (Figures 4M, 4N, and S3G-S3L). Although the most striking rearrangements of the plasma lipidome driven by *N*-oleoyl serinol were apparent in PC and ceramide lipid classes, there were several other notable alterations including altered circadian rhythms in fatty acid metabolites including 12-hydroxyeicosatetraenoic acid (12-HETE), 15-hydroperoxyeicosatetraenoic acid (15-HPETE), eicosapentaenoic acid (20:5), fatty acyl ester of hydroxy fatty acid 43:0 (FAHFA 43:0), phosphatidic acid 36:4 (PA 36:4), and diacylglycerol 50:14 (DG 50:14) (Figures S4M-S4R). In addition to clear alterations in the plasma lipidome, *N*-oleoyl serinol treatment also significantly altered circadian rhythms in carbohydrate-related and other metabolite classes including fructose, fructose-6-phosphate, α-ketoglutarate, citrulline, hydrocinnaminic acid, uric acid, and succinic acid (Figures 4O, S3S-S3X). In agreement with our findings under defined meal conditions (Figure 2), *N*-oleoyl serinol also reorganized the circadian rhythms of the gut microbiome (Figure S4). Even at the phylum level (Figure S4A-S4F) *N*-oleoyl serinol treatment reorganized the circadian oscillations of *Proteobacteria*, *Firmicutes*, and *Bacteroidota*. At the genus level (Figure S4G-S4R) there were clear alterations in the circadian oscillations of *Bifidobacterium*, *Acetitomaculum*, *Lactobacillus*, *Monoglobu*s, *Dubosiella*, *Enterorhabdus*, *GCA-900066575*, *family XIII UCG-001*, *Bacteroides*, *Odoribacter*, *Akkermansia*, and *Butyricicoccus* in *N*-oleoyl serinol treated mice. In parallel, the plasma levels of several gut microbe-associated metabolites were altered including trimethylamine (TMA), trimethylamine N-oxide (TMAO), γ-butyrobetaine, 4-hydroxyhippuric acid (4-OH-hippuric acid), indoxyl glucuronide, and serotonin (Figure S4S-S4X). Collectively, these studies show that *N*-oleoyl serinol can strikingly reorganize gene expression, metabolic hormones, and gut microbiome composition.

## DISCUSSION

Diet-microbe-host interactions clearly impact human disease^1–4,45–48^, but how our gut microbiota integrate dietary cues into meal-related signals to shape host phenotypes remains poorly understood. Here, we demonstrate that gut microbe-derived *N*-acyl amides exert broad effects on host hormonal control of metabolism after a meal. Given that *N*-acyl amides shape the postprandial levels of key metabolic hormones such as GLP-1 and insulin, this work has broad implications in diseases such as obesity, diabetes, fatty liver, and CVD. The major findings of the current study are: (1) *N*-oleoyl serinol increases postprandial insulin and leptin levels; (2) despite elevated postprandial insulin, hepatic lipogenic gene expression and cholesterol levels are reduced in *N*-oleoyl serinol-treated mice; (3) *N*-oleoyl serinol alters meal-related reorganization of the gut microbiome; (4) colonizing germ-free mice with gut bacteria harboring the *nas* gene alters metabolic hormones and metabolic gene expression during the fasting-to-feeding transition; and (5) elevated *N*-oleoyl serinol exerts strong pleiotropic effects on gene expression, metabolic hormones, gut microbiome composition, and circulating metabolites levels throughout a 24-hour circadian period. Collectively, our studies demonstrate that gut microbe-derived *N*-acyl amide lipids are underappreciated signals originating from the gut microbial endocrine organ that can bridge dietary intake to metabolic physiology.

Lipid signaling is a common means by which meal-related cues are relayed to the host to elicit diverse physiologic responses related to gastric motility, appetite, and anabolic versus catabolic metabolism^49–59^. For example, like the bacterially-derived *N*-acyl amides studied here, several host-derived acyl-ethanolamide and *N*-acylphosphatidylethanolamine (NAPE) lipids such as anandamide (AEA) oleoylethanolamide (OEA), and palmitoylethanolamide (PEA) can potently impact food intake, gastric motility, and metabolism following a meal^49–53^. In fact, drugs targeting host cannabinoid receptors sensing these amide lipids have profound effects on weight loss and diabetes resolution^54,55^. Although our understanding of diet-microbe-host interactions in lipid signaling is still in its infancy, it is likely that both bacterially and human derived lipids including acyl-amides, eicosanoids^56,57^, sphingolipids^26,58^, and likely many others could serve as key signaling intermediates in endocrine circuits. An emerging idea in the field states that gut bacteria can synthesize unique lipids that mimic host-derived lipid signals and engage host receptor systems to integrate physiological response^40^. It is also important to consider that some bacterially-derived lipids can be metabolically intertwined with host lipids via esterification into cell membranes^26,56–58^. Therefore, it is important to understand the integrated complexity of dietary substrate presentation, bacterial and host co-metabolism, and host receptor sensing and physiologic/pathophysiologic response to postprandial lipid signals. There is little doubt that amide-containing lipids are interesting therapeutic targets for metabolic disease^40,49–53^, but at this point essentially all drug discovery has been focused on host lipids. The present study argues for future drug discovery in the endocrine and metabolic space to consider both microbe-and host-targeted therapeutic strategies. For instance, could exogenous *N*-oleoyl serinol further improve efficacy and safety of GLP-1 receptor agonists? There are many unanswered questions regarding the role of the gut microbiome in the field of host endocrinology and metabolism, but it is reasonable to assume that there is untapped therapeutic potential targeting bacterial lipid metabolism to improve health in the human metaorganism.

### Limitations

First, it is important to note that we see slightly different results in metabolic physiology when *N*-oleoyl serinol is elevated via slow release pellets or via colonization with *nas*-encoding bacteria. This is likely due to the fact that mice harboring *nas*-encoding bacteria have elevated levels of several species of *N*-acyl amides on top of *N*-oleoyl serinol. Second, it is very likely that bacterially-derived *N*-acyl amides can be further metabolized by host enzymes and potentially incorporated into other complex lipids. Future studies using stable isotope or radioisotope tracers will be require to fully understand microbe-host co-metabolism of bacterially-derived *N*-acyl amides. It is interesting to note that a recent report has identified a bacterial fatty acid amide hydrolase enzyme that has the potential to degrade *N*-acyl amides^59^, but will require further investigation whether it is involved in the metabolic circuit studied here. Also, the only reported host receptor responsible for sensing bacterial *N*-oleoyl serinol is GPR119^40^. It is interesting to note that GPR119 has been previously identified as a therapeutic target in metabolic diseases including obesity and diabetes^60–62^. Therefore, future studies will need to investigate whether bacterially-derived *N*-acyl amides can impact the therapeutic efficacy of other GPR119 agonists. Finally, the present studies are exclusively done in mice. Whether similar diet-microbe-host signaling circuits occur in humans remains to be determined. Despite these limitations, this work demonstrates that *N*-acyl amides originating from the gut microbial endocrine organ can powerfully shape hormonal control of metabolism after a meal.

### Lead contact

Further information and requests for resources and reagents should be directed to and will be fulfilled by the lead contact, J. Mark Brown

### Materials availability

All the data and materials that support the findings of this study are available within the article and the *Online Supplement* or available from the lead author upon request.

### Data and code availability

All multi-omics datasets are currently being submitted to publicly-available data portals, and accession numbers will be provided upon acceptance of this work.

Code used for cosinor analyses:

1. Sachs, M. C. (2014) cosinor: Tools for estimating and predicting the cosinor model, version 1.1. R package: https://CRAN.R-project.org/package=cosinor.

Any additional information required to reanalyze the data reported in this paper is available from the lead contact (Dr. J. Mark Brown) upon request

## Acknowledgements

This work was supported in part by National Institutes of Health grants R01 DK130227 (J.M.B.), P01 HL147823 (S.L.H. and J.M.B.), P50 AA024333 (J.M.B.), RF1 NS133812 (J.M.B.), and an American Heart Association Postdoctoral Fellowship 24POST1178494 (S.D.). W.J.M. is supported by a Cleveland Clinic Global Center for Pathogen and Human Health Research Postdoctoral Fellowship provided by the Infection Biology Program of the Cleveland Clinic. The authors would like to thank investigators at the UC Davis West Coast Metabolomics center for acquiring some of the reported untargeted metabolomics data. The timsTof Pro2 instrument used for proteomic studies was purchased via an NIH shared instrument grant S10 OD030398 (B.W.).

## Author Contributions

Conceptualization, S.D., K.M., W.J.M., and J.M.B.; methodology, S.D., K.K.M., W.J.M., V.V., A.C.B., A.J.H., M.M., N.M., D.O., L.J.O., T.G., R.H., R.B., V.U., D.L., G.H., X.Y., N.S., M.D., A.M.H., B.W., and Z.W.; investigation, S.D., K.K.M., W.J.M., V.V., A.C.B., A.J.H., M.M., N.M., D.O., L.J.O., T.G., R.H., R.B., V.U., D.L., G.H., X.Y., N.S., M.D., B.W., and Z.W. validation, S.D., K.K.M., W.J.M., V.V., A.C.B., A.J.H., M.M., N.M., D.O., L.J.O., T.G., R.H., R.B., V.U., D.L., G.H., X.Y., N.S., M.D., B.W., and Z.W. formal analysis, S.D., K.K.M., W.J.M., T.G., N.S., Z.W., and J.M.B; writing – original draft, S.D., K.K.M., W.J.M., and J.M.B.; writing – reviewing and editing, S.D., K.K.M., W.J.M., V.V., A.C.B., A.J.H., M.M., N.M., D.O., L.J.O., T.G., R.H., R.B., V.U., D.L., G.H., X.Y., N.S., M.D., A.M.H., B.W., and Z.W., S.L.H, and J.M.B.; funding acquisition, S.L.H. and J.M.B.; supervision, J.M.B.

## ABBREVIATIONS USED

*Acaca*: acetyl-CoA carboxylase alpha
*Acly*: ATP-citrate lyase
AEA: arachidonoylethanolamide
*Bhlhe40*: basic helix-loop-helix family member e40
*Cpt1α*: carnitine palmitoyltransferase 1 alpha
*Cry2*: cryptochrome 2
CVD: cardiovascular disease
*Fasn*: fatty acid synthase
*Fgf21*: fibroblast growth factor 21
G6pd: glucose-6-phosphate dehydrogenase
ψBB: gamma butyrobetaine
GLP-1: glucagon-like peptide 1
GPR119: G protein-coupled receptor 119
*Hmgcr*: 3-hydroxy-3-methylglutaryl-CoA reductase
IL-6: interleukin 6
Irs2: insulin receptor substrate 2
MCP-1: monocyte chemoattractant protein 1
*N*-AS: *N*-arachidonoyl serinol
*N*-OS: *N*-oleoyl serinol
*N*-PS: *N*-palmitoyl serinol
*Nr1d1*: nuclear receptor subfamily 1 group D member 1 or REV-ERBalpha
OEA: oleoylethanolamide
PC: phosphatidylcholine
*Pck1*: phosphoenolpyruvate carboxykinase
PEA: palmitoylethanolamide
*Pgc1α*: peroxisome proliferator-activated receptor gamma coactivator 1-alpha
*Pnpla3*: patatin-like phospholipase domain-containing protein 3
PYY: peptide YY
SEA: stearoylethanolamide
*Soat1*: sterol-O-acyltransferase 1
*Soat2*: sterol-O-acyltransferase 2
*Srebf1*: sterol regulatory element-binding transcription factor 1
*Srebf2*: sterol regulatory element-binding transcription factor 2
TMA: trimethylamine
TMAO: trimethylamine oxide
TNFα: tumor necrosis factor α
*Ucp1*: uncoupling protein 1
WAT: white adipose tissue
ZT: zeitgeber time

## HIGHLIGHTS

- Gut bacterial-derived *N*-oleoyl serinol alters meal-related hormone levels.
- *N*-oleoyl serinol dampens postprandial lipogenic response in the liver.
- Gnotobiotic mice harboring the *nas* gene exhibit altered metabolic homeostasis in the host.
- *N*-oleoyl serinol shapes meal- and circadian-driven changes of microbiota and host metabolism.

### eTOC BLURB

Diet-microbe-host interactions clearly impact human disease, but how gut microbiota integrate dietary cues into meal-related signals to shape host phenotypes remains poorly understood. Dutta *et al.* now show that gut microbe-produced N-acyl serinols broadly alter meal- and circadian-driven alterations in gene/protein expression, hormones, microbiota, and metabolite levels in the host.

## METHODS

### Chemical Synthesis of *N*-Palmitoyl Serinol and *N*-Oleoyl Serinol

The small molecules *N*-Palmitoyl Serinol (*N*-PS) and *N*-Oleoyl Serinol (*N*-OS) were synthesized and structurally characterized as outlined below using multinuclear NMR and mass spectrometry. ^1^H and ^13^C NMR spectra were acquired on a JEOL-ECZ 400S spectrometer equipped with a 5 mm ROYALProbe HX probe operating at a nominal 400 MHz. Chemical shifts are reported in parts per million (ppm) and referenced to the DMSO-D6 residual solvent signal at 2.50 ppm. *N*-PS was prepared by activating 15g of Palmitic acid to the *N*-Hydroxysuccinimide ester using *N*-Hydroxysuccinimide (NHS) and 1,3-Dicyclohexylcarbodiimide (DCC) in a dry dichloromethane and tetrahydrofuran mixture, followed by amidation to 2-amino-1,3-propanediol pre-dissolved in ethanol. The resulting crude product was isolated by flash chromatography using (1:9 / methanol:chloroform) and recrystallized from warm methanol to achieve final purity (HPLC, 900:100:1 Hexane:Isopropanol:Acetic Acid, Silica 4.6 x 250 mm, 5 um particle, 1 mL/min at UV 210 nm: 99.2% area, 14.28g, 43.34mmol, 74% yield). MS corroborated the expected cation mass (MS (ESI): m/z ([M+H]^+^) calculated for C_19_H_40_NO_3_^+^ ([M+H]^+^] 330.3, found 330.2). ^1^H and ^13^C NMRs were obtained and the ^1^H NMR was consistent with literature^63^ and with expected values. ^1^H NMR (400 MHz, DMSO-d_6_): δ 7.42 (br d, *J* = 8.0 Hz, 1H, -C(O)N<u>H</u>-CH-), 4.56 (t, *J* = 5.6 Hz, 2H, -CH_2_-O<u>H</u>), 3.68 (dquin, *J* = 8.0, 5.5 Hz, 1H, -C(O)NH-C<u>H</u>(CH_2_-)_2_), 3.37 (t, *J* = 5.5 Hz, 4H, -CH(C<u>H</u>_2_-OH)_2_), 2.05 (t, *J* = 7.5 Hz, 2H, -CH_2_-C<u>H</u>_2_-C(O)NH-), 1.46 (quin, *J* = 6.9 Hz, 2H, -C<u>H</u>_2_-CH_2_-C(O)NH-), 1.23 (m, 24H, (-C<u>H</u>_2_-)_12_), 0.85 (t, *J* = 6.6 Hz, 3H, -CH_2_-C<u>H</u>_3_); ^13^C NMR (100 MHz, DMSO-d_6_): δ 172.09 (-CH_2_-<u>C</u>(O)NH-CH-), 60.20 (2C, -CH(<u>C</u>H_2_-OH)_2_), 52.72 (-C(O)NH-<u>C</u>H(CH_2_-OH)_2_), 35.42 (-CH_2_-<u>C</u>H_2_-C(O)NH-), 31.30 (CH_3_CH_2_-<u>C</u>H_2_-CH_2_-), 29.5-28.5 (10C, peaks at 29.06, 29.05, 29.03, 28.98, 28.84, 28.72, 28.67 (-CH_2_-(<u>C</u>H_2_)_10_-CH_2_)), 25.34 (-CH_2_-<u>C</u>H_2_-CH_2_C(O)NH-), 22.11 (CH_3_-<u>C</u>H_2_-CH_2_-), 13.97 (<u>C</u>H_3_-CH_2_-) *N*-OS was prepared using *N*-Hydroxysuccinimide (NHS) and *N*-(3-Dimethylaminopropyl)-N′-ethylcarbodiimide hydrochloride (EDC) to activate 20g of Oleic acid to the *N*-Hydroxysuccinimide ester, which was isolated via liquid/liquid extraction with dichloromethane (DCM) from water and brine to remove byproducts, followed by rigorous drying. Amidation to 2-amino-1,3-propanediol in DCM and ethanol produced crude product that was isolated via liquid/liquid extraction with ethyl acetate from water and brine, followed by rigorous drying to afford pure final *N*-OS (HPLC, 915:84:1 Hexane:Isopropanol:Acetic Acid, Silica 4.6 x 250 mm, 5 um particle, 2 mL/min at UV 210 nm: 100.0% area, 21.2g, 59.6mmol, 84.3% yield). MS corroborated the expected cation mass (MS (APCI): m/z ([M+H]^+^) calculated for C_21_H_42_NO_3_^+^ ([M+H]^+^] 356.3, found 356.3). ^1^H and ^13^C NMRs were obtained and the ^1^H NMR was consistent with literature^64^ and with expected values. ^1^H NMR (400 MHz, DMSO-d_6_): δ 7.43 (br d, *J* = 8.0 Hz, 1H, -C(O)N<u>H</u>-CH-), 5.42 – 5.23 (m, 2H, -CH_2_-C<u>H</u>=C<u>H</u>-CH_2_), 4.57 (t, *J* = 5.6 Hz, 2H, -CH_2_-O<u>H</u>), 3.68 (dquin, *J* = 8.0, 5.5 Hz, 1H-C(O)NH-C<u>H</u>(CH_2_-)_2_), 3.37 (t, *J* = 5.5 Hz, 4H, -CH(C<u>H</u>_2_-OH)_2_), 2.05 (t, *J* = 7.5 Hz, 2H, -CH_2_-C<u>H</u>_2_-C(O)NH-), 1.98 (q, *J* = 6.3 Hz, 4H, -CH_2_-C<u>H</u>_2_-CH=), 1.46 (quin, *J* = 7.2 Hz, 2H, -C<u>H</u>_2_-CH_2_-C(O)NH-), 1.36 – 1.13 (m, 20H, (-C<u>H</u>_2_-)_10_), 0.85 (t, *J* = 6.9 Hz, 3H, -CH_2_-C<u>H</u>_3_); ^13^C NMR (100 MHz, DMSO-d_6_): δ 172.07(-CH_2_-<u>C</u>(O)NH-CH-), 129.65 (s, 2C, -CH_2_-<u>C</u>H=<u>C</u>H-CH_2_-), 60.20 (s, 2C, -CH(<u>C</u>H_2_-OH)_2_), 52.71 (-C(O)NH-<u>C</u>H(CH_2_-OH)_2_), 35.41 (-CH_2_-<u>C</u>H_2_-C(O)NH-), 31.28 (CH_3_CH_2_-<u>C</u>H_2_-CH_2_-), 29.2-28.5 (8C, peaks at 29.14, 29.10, 28.83, 28.72, 28.69, 28.58 (-CH_2_-<u>C</u>H_2_-CH_2_-)), 26.61 (CH_2_-<u>C</u>H_2_-CH=), 26.57 (CH_2_-<u>C</u>H_2_-CH=), 25.34 (-CH_2_-<u>C</u>H_2_-CH_2_C(O)NH-), 22.10 (CH_3_-<u>C</u>H_2_-CH_2_-), 13.96 (<u>C</u>H_3_-CH_2_-)

### Engineering of *E. coli* to Stably Express the *nas* Gene

To specifically understand the role of gut microbe-derived N-acyl amides in host metabolic physiology we cloned the bacterial *nas* gene from *Gemella haemolysans* M34^20^ and stably expressed it in a mouse derived *E. coli* strain (MP1)^65^. *E. coli* MP1 was a kind gift from Dr. Mark Joulian at University of Pennsylvania. DNA sequence corresponding to *Gemella haemolysans nas* gene was PCR amplified and placed downstream a synthetic medium strength promotor (PJ23114, http://parts.igem.org/Part:BBa_J23114) and an optimized ribosomal binding site (RBS). rrnB operon terminator was placed downstream the expression cassette. The casstte was then flanked on both sides by ∼ 1kb homologous recombination arms so that the cloned nas gene is placed between Kch (<u>WPV03328.1)</u> and YciY (WPV03329.1) genes on E*. coli* MP1 chromosome. The whole PCR product was then ligated to a suicide plasmid using In-Fusion® HD Cloning kit (Clontech) and transformed into *E. coli* S17 λpir. The suicide plasmid contained kanamycin resistance cassette*, sacB* as counter selection marker, R6K replication origin, and RP4-oriT. In parallel, *E. coli* MP1 was trasnofrmed with a temperature sensitive plasmid coding for ampicillin resistance. The suicide plasmid was transferred from *E. coli* S17 λpir to amicillin-resistant *E. coli* MP1 through conjugation. Resulted *E. coli* MP1 merodiploid conjugants were plated on LB agar plates containing kanamycin at 50 ng/µl and carbenicillin at 200 ng/µl. One *E. coli* MP1 merodiploid mutant was selected and re-streaked on LSW-Sucrose agar plate containing 10% sucrose^66^. Screening for knockin mutants was done through PCR, and the correct knockin was re-streaked, and confirmed for loss of conjugated plasmid through DNA sequencing. To cure constructed *E. coli* MP1 mutant from the ampicillin-resistance plasmid, they were cultured at 40 °C and screened for loss of ampicillin resistance.

### Animal Studies (Mice)

To test the long-term consequence of *N*-oleoyl serinol treatment, slow-releasing pellets containing either placebo or 4.9 mg *N*-oleoyl serinol (350 μg/day) were implanted subcutaneously under isoflurane anesthesia in conventionally housed C57BL/6J mice purchased from Jackson Laboratories (Bar Harbor, MN, USA). Pellets were custom-made (by Innovative Research of America, Sarasota, FL) to release either placebo or *N*-oleoyl serinol continuously for 14 days. Briefly, a small (5 mm) incision was made at the nape of the neck, the pellet was inserted and the incision closed with sutures. For the fasting and re-feeding studies 8 week old male C57BL/6J mice were procured from Jackson Laboratories, and were kept in the mouse housing facility at 25°C with continuous access to food and water at the Cleveland Clinic Biological Resource Unit. The mice were fed a standard chow diet (Teklad Global Diet 2018S, Inotiv). After 4 weeks, these mice underwent a surgery where small 14-day slow release placebo or *N*-oleoyl serinol pellets (350 µg/day; Innovative Reserach of America, FL) were subcutaneously implanted as previously described^18,41^. Mice were thereafter kept on *ad libitum* conditions for 7 days to allow for N-oleoyl serinol levels to reach steady state and then were necropsied in the fed state (*ad libitum*), or 12 hour fasted state (fasted from 2200 h to 1000 h), or refed state (12 hours of fasting (2200 h to 1000 h), followed by 3 hours of refeeding (1000 h to 1300 h). For the circadian *N*-oleoyl serinol pellet implant studies, 8-week-old male C57BL/6J mice were purchased from Jackson Laboratories and placed on defined chow diet (Envigo diet # TD.130104) one week before light/dark acclimation. Mice were transferred to a room equipped with 12:12 light-dark cycling for one week before pellet implantation surgeries were performed. Pellet surgeries were performed during daylight hours as described above, and mice were kept in a room equipped with 12:12 light:dark cycling for one week before sacrifice. One week after the pellet implantation, plasma and tissue collection were performed every 4 hours (Zeitgeber times 2, 6, 10, 14, 18, and 22) over a 24-hour period. All dark cycle necropsies were performed under red light conditions. For the gnotobiotic studies, 23-28-week-old germ-free C57BL/6N mice were generated within the Cleveland Clinic Gnotobiotic Core facility. Germ-free recipient mice were given a single oral gavage of 200 μL wild type or *nas*-expressing *E. coli* in sterile PBS + 20% glycerol. Gnotobiotic colonized mice were maintained for three weeks using the Allentown Sentry SPP Cage System (Allentown, NJ) on a 14 hour:10 hour light: dark cycle. Mice in the fed group had ad libitum access to food and were necropsied first. The fasted group were fasted overnight (∼16 hours) prior to necropsy. For all mouse studies blood was collected in EDTA coated tubes by cardiac puncture at necropsy and were immediately processed for plasma isolation by centrifuging at 5500 g, 4°C for 5 mins. The translucent upper layer of plasma was transferred to a fresh cryovial and stored at -80°C until further downstream processing. The liver and gWAT tissues were collected at necropsy, weighed and a part of each tissue were processed for histology, while the remaining part was snap frozen in liquid nitrogen. The gastrointestinal tract was sectioned into duodenum, jejunum, ileum, cecum and colon, and were snap frozen. All tissues were stored at -80°C until further downstream processing. All mice were maintained in an Association for the Assessment and Accreditation of Laboratory Animal Care, International-approved animal facility, and all experimental protocols were approved by the Institutional Animal Care and use Committee of the Cleveland Clinic (Approved IACUC protocol numbers are #2019-2207, #00002867, and #00003407).

### RNA and Realtime PCR Methods

RNA isolation and quantitative polymerase chain reaction (qPCR) was conducted using methods previously described with minor modifications^67–69^. To extract RNA, the Monarch Total RNA MiniPrep Kit (New England BioLabs, Inc., Cat# T2010S) was employed, and cDNA was synthesized using Quanta Bio qScript cDNA Supermix (Cat# 95048-100) in a c1000 Touch Thermal Cycler (Biorad, Hercules, CA). For the fed/fasted/refed pellet implantation study and the *nas*-expressing *E. coli* gnotobiotic study, 800 ng of cDNA was utilized for Realtime PCR on a 96-well plate, using Applied Biosystem’s Fast SYBR™ Green Master Mix (Cat# 4385618) in a Applied Biosystems™ StepOne™ Real-Time PCR System (Applied Biosystems Corp., Waltham, MA). For the circadian study with *N*-OS pellet treatment, gene expression analyses was performed on 384-well plate, Applied Biosystem’s Fast SYBR™ Green Master Mix in either a Roche LightCycler 480 I1 or Biorad CFX Opus 384 well PCR machine. The levels of induced mRNAs were standardized by the ΔΔCT method and were normalized to cyclophilin. The specificity of the primers was verified by analyzing the melting curves of the PCR products. Primer information is available in the Supplemental table S1.

### Quantification of *N*-Acyl Amide Lipids By Liquid Chromatography Tandem Mass Spectrometry (LC-MS/MS)

For quantifying circulating levels of *N*-acyl amides in peripheral or portal plasma, 20 μl of plasma was transferred to heat-sterilized 12 x 75 mm glass tubes (Fisherbrand™ Round Bottom Disposable Borosilicate Glass Tubes, Fisher Scientific; Cat# 14-961-26). To the plasma, 10 μl of the internal standard mix in methanol (containing C_18:1_EA-d5, C_16:0_EA-d5, C_20:4_EA-d5, C_16:0_Ser-d5 and C_18:1_Ser-d5, at a concentration of 1 μM each) and 380 μl of LC-MS grade water were added, and mixed by vortexing for ∼1 min. To the mixture, 1 ml of solution A (containing 2-propanol, hexanes and acetic acid with a ratio of 140: 60: 0.23, v/v/v) was added and vortexed well. Thereafter, an additional 1 ml of hexanes was added to the tube and the contents were mixed well by vortexing. Hexanes solubilizes the *N*-acyl amides and separates them from the bottom aqueous layer. For proper phase separation, each sample was centrifuged at 2500 g at 4°C for 5 mins. The upper solvent layer of hexanes containing *N*-acyl amides was then transferred to a fresh glass tube, and the contents were dried under a constant flow of N_2_ gas. All samples were stored at this stage at -20°C before the MS run. 20 ul of standards (ranging from 5 – 500 nM) were also processed in parallel with plasma samples.

For quantifying the level of *N*-acyl amides in liver tissues, ∼25 mg of liver tissue were homogenized in 500 μl (20 times the tissue weight, 1 mg tissue corresponds to 1 μl in volume) of the internal standard mix in methanol (containing C_18:1_EA-d5, C_16:0_EA-d5, C_20:4_EA-d5, C_16:0_Ser-d5 and C_18:1_Ser-d5 at a concentration of 50 nM each) in Eppendorf Safe-Lock Tubes (2.0 mL) using a Qiagen Tissue Lyser II (Qiagen, Germany). To remove any residual debris, the tissue lysates were centrifuged at 20,000 g at 4°C for 5 mins. The clear supernatants were then transferred to heat-sterilized 12 x 75 mm glass tubes (Fisherbrand™ Round Bottom Disposable Borosilicate Glass Tubes, Fisher Scientific; Cat# 14-961-26), and the contents were dried using a speed vacuum. Thereafter, the dried lysates were resuspended in 400 μl of LC-MS grade water and 1 ml of 2-propanol : hexanes : 2M acetic acid solution (40:10:1, v/v/v), and vortexed for 1 min for proper mixing. Another 1 ml of hexanes was added to the mixture and vortexed for another 1 min. Then, the samples were centrifuged at 2500 g at 4°C for 5 mins and the upper solvent layer of hexanes containing *N*-acyl amides was then transferred to a fresh glass tube. The contents of the tube were dried under a constant flow of N_2_ gas and all samples were stored at -20°C before the MS run. 20 ul of standards (ranging from 50 – 1000 nM) were also processed in parallel with plasma samples.

The well-prepared MS sample was resuspended in methanol followed by spin-down. Supernatants (10 μl) were analyzed by injection onto a Kinetex™ C18 HPLC Column (2.6 μm particle size, L × I.D. 50 mm × 2.1 mm, #00B-4462-AN, Phenomenx) at a flow rate of 0.35 ml min^−1^ using a 2 Shimadazu LC-20AD Nexera CL pump system, SIL-30AC MP CL autosampler interfaced with an Shimadzu 8050 mass spectrometer. A discontinuous gradient was generated to resolve the analytes by mixing solvent A (0.1% formic acid in water) with solvent B (0.1% formic acid in acetonitrile) at different ratios starting from 50% B for 2 minutes, then linearly to 83% B over 4 min, then to 100% B over 0.2 min and hold for 3 min, and then back to 50% B. Amide lipids and their respective isotope labeled internal standards were monitored using electrospray ionization in positive-ion mode with multiple reaction monitoring (MRM) of precursor and characteristic product-ion transitions of *m/z* 300.3 →62.1, 305.3 →62.1, 326.3→ 62.1, 328.3→ 62.1, 348.3→ 62.1, 352.3→ 66.1, 330.3→ 312.4, 335.3→ 317.3, 356.3→ 92.1, 361.4→ 97.1, 378.3→ 92.1, and 269.3→ 251.35 amu, for palmitoylethanolamide, [^2^H_5_] palmitoylethanolamide, oleoylethanolamide, [^2^H_2_] oleoylethanolamide, arachidonoylethanolamide, [^2^H_4_] arachidonoylethanolamide, palmitoylserinol, [^2^H_5_] palmitoylserinol, oleoylserinol, [^2^H_5_] oleoylserinol, arachidonoylserinol and cis-9,10-Methylenehexadecanoic acid (CMHA), respectively. The parameters for the ion monitoring were optimized individually. Nitrogen (99.95% purity) was used as the source and argon was used as collision gas. Various concentrations of standard mix were mixed with internal standard mix undergoing the same sample processing steps were used to prepare the calibration curves for quantification of amide lipids and CMHA. [^2^H_5_] oleoylserinol was used as internal standard for calibration of arachidonoylserinol and CMHA. Stearoylethanolamide was monitored in the same characteristic product-ion transition as [^2^H_2_] oleoylethanolamide but with different retention time from [^2^H_2_] oleoylethanolamide, and [^2^H_2_] oleoylethanolamide was used as internal standard for calibration of stearoylethanolamide.

### Quantification of Other Gut Microbe-Derived Metabolites Using LC-MS/MS

For this study we wanted to broadly understand how gut microbe-derived metabolites originating from diverse dietary substrates are altered by *N*-acyl amide administration. Therefore, we used a stable-isotope-dilution liquid chromatography tandem mass spectrometry (LC-MS/MS) methods for the quantitative analysis of diverse bacterially derived metabolites that originate from dietary micronutrients such as choline and carnitine, as well as aromatic amino acids including phenylalanine, tyrosine and tryptophan. A total of 20 μL of mouse plasma was used for extraction of metabolites using the method described by Nemet and colleagues^21^. Briefly, 80 μL of isotope labeled internal standard mix was added to 20 μL of plasma, vortexed for 1 min, followed by centrifugation at 20,000 g, in 4 °C for 10 min. A total of 80 μL of supernatant was transferred to a MS vial with plastic insert and 0.5 μL was used for LC-MS. The plasma concentration of gut microbe-associated metabolites originating from aromatic amino acids was recently described by Nemet and colleagues^21^. Trimethylamine (TMA) related metabolites were measured as previously descibed by Wang and colleagues^28^.

### Untargeted Lipidomics Using Liquid Chromatography Tandem Mass Spectrometry (LC-MS/MS)

Samples extracted using Matyash extraction procedure which includes MTBE, MeOH, and H2O^70^. The organic (upper) phase was dried down and submitted for resuspension and injection onto the LC. Samples are resuspended with 110 μL of a solution of 9:1 methanol: toluene and 50 ng/mL CUDA. This is then shaken for 20 seconds, sonicated for 5 minutes at room temperature, and then centrifuged for 2 minutes at 16100 rcf. The samples are then aliquoted into three parts. 33 μL are aliquoted into a vial with a 50 μL glass insert for positive and negative mode lipidomics. The last part is aliquoted into an eppendorf tube to be used as a pool. The samples are then loaded up on an Agilent 1290 Infinity LC stack. The positive mode was run on an Agilent 6530 with a scan range of m/z 120-1200 Da with an acquisition speed of 2 spectra/s. Positive mode has between 0.5 and 2 μL injected onto an Acquity Premier BEH C18 1.7 µm, 2.1 x 50 mm Column. The gradient used is 0 min 15% (B), 0.75 min 30% (B), 0.98 min 48% (B), 4.00 min 82% (B), 4.13-4.50 min 99% (B), 4.58-5.50 min 15% (B) with a flow rate of 0.8 mL/min.

The other sample aliquot was run in negative mode, which was run on Agilent 1290 Infinity LC stack, and injected on the same column, with the same gradient and using an Agilent 6546 QTOF mass spec. The acquisition rate was 2 spectra/s with a scan range of m/z 60-1200 Da. The mass resolution for the Agilent 6530 is 10,000 for ESI (+) and 30,000 for ESI (-) for the Agilent 6546.

### Untargeted Metabolomics Using Gas Chromatography-Time of Flight (TOF)-Mass Spectrometry

Samples extracted using Matyash extraction procedure which includes MTBE, MeOH, and H2O. The aqueous (bottom) phase was dried down and submitted to derivatization for GC. Samples are shaken at 30C for 1.5 hours. Then 91 uL of MSTFA + FAMEs to each sample and they are shaken at 37C for 0.5 hours to finish derivatization. Samples are then vialed, capped, and injected onto the instrument. We use a 7890A GC coupled with a LECO TOF. 0.5 uL of derivatized sample is injected using a splitless method onto a RESTEK RTX-5SIL MS column with an Intergra-Guard at 275C with a helium flow of 1 mL/min. The GC oven is set to hold at 50C for 1 min then ramp to 20C/min to 330C and then hold for 5 min. The transfer line is set to 280C while the EI ion source is set to 250C. The Mass spec parameters collect data from 85m/z to 500m/z at an acquisition rate of 17 spectra/sec.

### Quantification of Plasma and Hepatic Lipids

Hepatic lipids were extracted, and both plasma and hepatic lipids (triglycerides, phospholipids, total cholesterol and free cholesterol) were quantified using commercially available enzymatic assay kits from Fujifilm and Thermo Scientific (details in Supplemental table S1) as previously described^67–69^.

### Global Proteomics of Mouse Liver

Mouse liver tissue samples were suspended in 150 μL 8M urea Tris-HCl pH8 lysis buffer with freshly added protease inhibitor cocktail. Samples were homogenized by ultrasonication 10s x 3 with 10s between runs. Homogenized samples were centrifuged at 15000 kg for 15 minutes, and the supernatants were transferred to new 1.5 mL tubes. Protein concentrations of the samples were determined by BCA. A 20 μg protein aliquot from each of the samples were reduced by dithiothreitol, alkylated by iodoacetamide and precipitated by cold acetone (−20°C) overnight. Samples were centrifuged at 12000 g for 15 minutes at 0 °C, and the supernatants were removed. Protein pellets were air dried for 30 minutes and dissolved in 40 μL 50mM tri-ethyl ammonium bicarbonate (TEAB) with 0.3 μg sequencing grade trypsin and incubated at room temperature overnight. Five μg of peptides were taken from each sample and dried in a SpeedVac and reconstituted in 25 µl 0.1% formic acid. Ten µl sample was mixed with 10 µl 0.5x concentration iRT standards and the samples are ready for LCMS analysis. The tissue samples were analyzed using a Data Independent Acquisition (DIA) strategy. For these experiments, a NanoElute HPLC was coupled to a TimsTof Pro2 Q-Tof mass spectrometry system (Bruker Daltonik GmbH, Bremen). The HPLC was equipped with a Bruker 15 cm x 75 µm id C18 ReproSil AQ, 1.9 μm, 120 Å reversed-phase capillary chromatography column. Peptide separation was performed using a 50 minute gradient from 2 to 35% mobile phase B where mobile phase A was 0.1% formic acid and mobile phase B was acetonitrile/0.1% formic acid at a flow rate of 0.3 μL/min. To build a spectral library, 10 μg from each of the digested samples were pooled. The pooled sample was desalted using a Waters C18 Sep-Pak cartridge. The desalted pooled sample was off-line fractionated into 16 fractions using a high pH reversed phase HPLC method on a Waters XBridge BEH C18 2.1 mm X 150 mm, 3.5 μm, 130Å chromatographic column using a 48-minute from 2 to 90% mobile phase B where mobile phase A was 0.1% formic acid 10 mM ammonium formate in water and mobile phase B was 90% acetonitrile/0.1% formic acid 10 mM ammonium formate in water at a flow rate of 0.25 mL/min. Fractions were dried in a SpeedVac and reconstituted in 50 μl 0.1% formic acid each. 10 μl from each fraction was mixed with 10 μl 0.5x concentration iRT standards and the samples were ready for LCMS analysis. For spectral library generation, the fractionated samples were analyzed using a Parallel Accumulation–Serial Fragmentation (PASEF) data dependent acquisition (DDA) method. This method utilized MS1 scans for the identification of precursor ions followed by 10 PASEF MS/MS scans. The tims-MS survey scan was acquired between 0.60 and 1.43 Vs/cm2 and 100–1,700 m/z with a ramp time of 100 ms. The total cycle time for the PASEF scans was 1.17 seconds and the MS/MS experiments were performed with collision energies between 20 eV (0.6 Vs/cm2) and 59 eV (1.6 Vs/cm2). Precursors with 2–5 charges were selected with the target value set to 20,000 a.u and intensity threshold to 2,500 a.u. Precursors were dynamically excluded for 0.4 minutes.

For protein quantitation, each sample was analyzed using a PASEF data independent acquisition (DIA) method which includes a MS1 full scan followed by MS/MS scans of 32 fixed mass windows between 400 m/z and 1201 m/z. The TIMS-MS survey scan was acquired between 0.60 and 1.43 Vs/cm2 and 100–1,700 m/z with a ramp time of 100 ms. The total cycle time for the PASEF scans was 1.8 seconds and the MS/MS experiments were performed with collision energies between 20 eV (0.85 Vs/cm2) and 59 eV (1.3 Vs/cm2). The spectral library was generated using Pulsar integrated in Spectronaut V17 to search the DDA LC-MS data against the mouse SwissProt protein sequence database (Downloaded on 6-14-2021 with 17050 Entries). Oxidation of Met and protein N-terminal Acetylation were set as variable modifications and carbamidomethylation of Cys was set as fixed modification. The false discovery rate (FDR) of protein, peptide, and peptide spectral match (PSM) were all set to 1% of the identifications from a decoy database generated by reversing the amino acid sequences of the mouse SwissProt protein database. The iRT Reference Strategy was set to Deep learning assisted iRT regression. The DIA data of the samples were searched using Spectronaut V17 software against the spectral library and the mouse SwissProt protein sequence database. The relative abundance of the proteins in these samples was determined using a label free quantitation (LFQ) approach. Protein quantities were expressed as normalized intensities by Spectronaut V17. These are based on the sum of the (raw) intensities of the MS/MS product ions of their peptide precursor peaks that are normalized on total peptide quantities of each sample to ensure that profiles of the abundances across samples accurately reflect the relative amounts of the proteins.

### Plasma Hormone and Cytokine Quantification

Plasma hormones and cytokine levels were quantified using U-PLEX (product # K15297K) assays per the manufacturer’s instructions (Meso Scale Diagnostics, Rockville, Maryland, USA).

### 16S#rRNA gene amplicon sequencing and bioinformatics

16S rRNA gene amplicon sequencing and bioinformatics analysis were performed using our published methods^68^. Briefly, raw 16S amplicon sequence and metadata, were *demultiplexed using split_libraries_fastq.*py script implemented *in QIIME2*^71^. Demultiplexed fastq file was split into sample-specific fastq files using split_sequence_file_on_sample_ids.py script from QIIME2. Individual fastq files without non-biological nucleotides were processed using Divisive Amplicon Denoising Algorithm (DADA) pipeline^72^.The output of the dada2 pipeline [feature table of amplicon sequence variants (an ASV table)] was processed for alpha and beta diversity analysis using *phyloseq*^73^, and microbiomeSeq (http://www.github.com/umerijaz/microbiomeSeq) packages in R. We analyzed variance (ANOVA) among sample categories while measuring the of α-diversity measures using plot_anova_diversity function in *microbiomeSeq* package. Permutational multivariate analysis of variance (PERMANOVA) with 999 permutations was performed on all principal coordinates obtained during CCA with the *ordination* function of the *microbiomeSeq* package. Pairwise correlation was performed between the microbiome (genera) and metabolomics (metabolites) data was performed using the microbiomeSeq package.

### 16S Statistical Analysis

Differential abundance analysis was performed using the random-forest algorithm, implemented in the DAtest package (https://github.com/Russel88/DAtest/wiki/usage#typical-workflow).

Briefly, differentially abundant methods were compared with False Discovery Rate (FDR), Area Under the (Receiver Operator) Curve (AUC), Empirical power (Power), and False Positive Rate (FPR). Based on the DAtest’s benchmarking, we selected lefseq and anova as the methods of choice to perform differential abundance analysis. We assessed the statistical significance (*P* < 0.05) throughout, and whenever necessary, we adjusted *P*-values for multiple comparisons according to the Benjamini and Hochberg method to control False Discovery Rate^74^. Linear regression (parametric test), and Wilcoxon (Non-parametric) test were performed on genera and ASVs abundances against metadata variables using their base functions in R (version 4.1.2; R Core Team, 2021).

### Data Analyses for Circadian Rhythmicity (Cosinor Analyses)

A single cosinor analysis was performed as previously described^68^. Briefly, a cosinor analysis was performed on each sample using the equation for cosinor fit as follows:
Y(t)=M+Acos(2θ/τ+ϕ)
where M is the MESOR (midline statistic of rhythm, a rhythm adjusted mean), A is the amplitude (a measure of half the extent of the variation within the cycle), Φ is the acrophase (a measure of the time of overall highest value), and τ is the period. The fit of the model was determined by the residuals of the fitted wave. After a single cosinor fit for all samples, linearized parameters were then averaged across all samples allowing for calculation of delineralized parameters for the population mean. A 24 hour period was used for all analysis. Comparison of population MESOR, amplitude, and acrophase was performed as previously described^75^. Comparisons are based on F-ratios with degrees of freedom representing the number of populations and total number of subjects. All analyses were done in R v.4.0.2 using the cosinor and cosinor2 packages^75–78^ (Supplemental Table S3).

### Standard Statistical Analyses

All data are expressed as the mean ± standard error of the mean (S.E.M.). Other than 16S and circadian cosinor analyses, which are described above, all data were analyzed using one-way of variance followed by Student’s t tests for post hoc analysis. Differences were considered significant at p <0.05. All group means comparisons were performed using JMP statistical discovery software (version 17.0.0; SAS Institute).

