## Supplementary material for "Gut Microbe-Derived *N*-Acyl Serinol Lipids Shape Host Postprandial Metabolic Homeostasis": Online supplement-Figures S1-S4

**Journal, Volume XX**

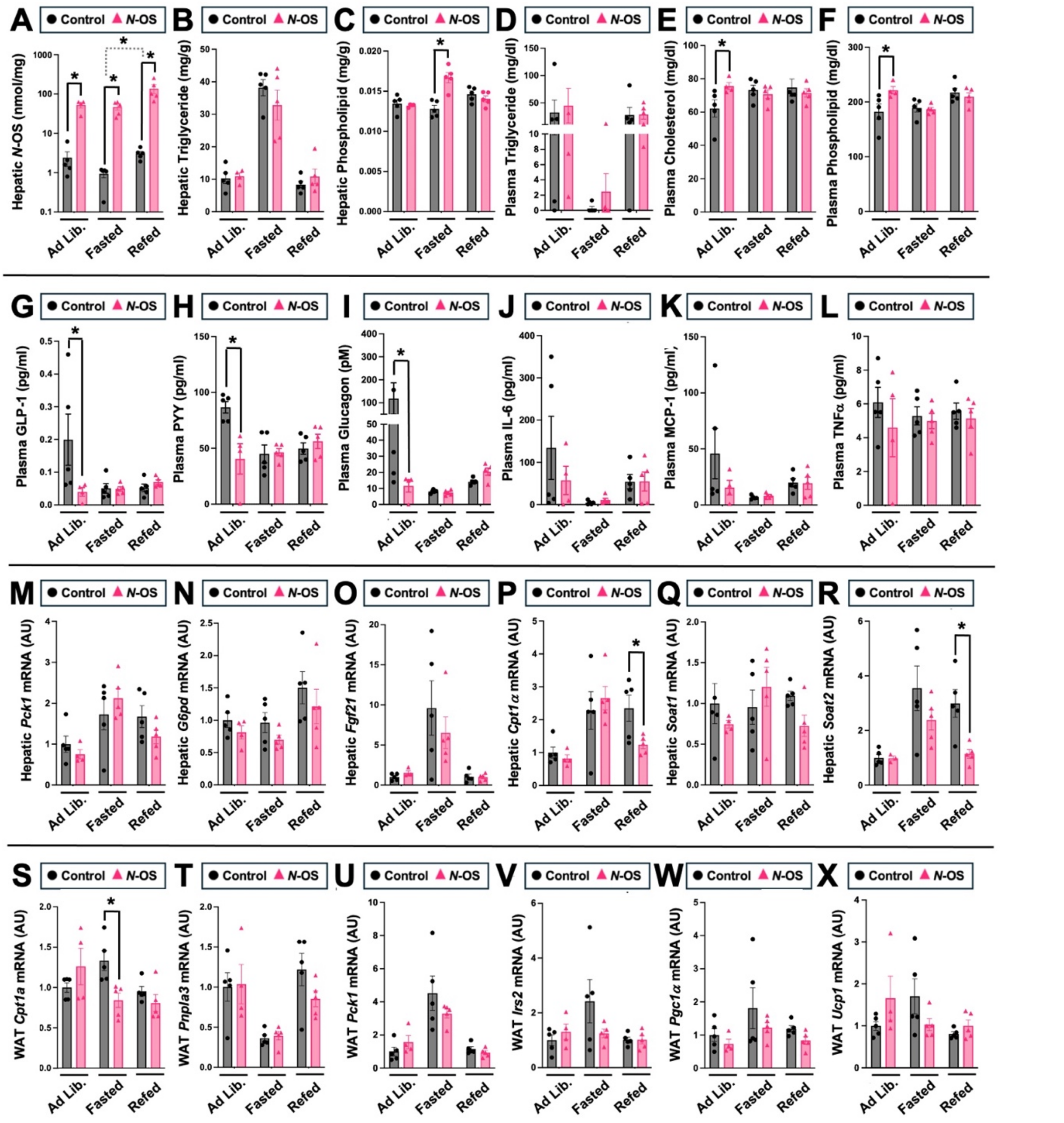


**Figure S1. *N*-Oleoyl Serinol Alters Multiple Aspects of Postprandial Hormonal and Metabolic Homeostasis (*Related to Figure 1*).** Male chow-fed C57BL/6J mice were implanted with slow-release pellets containing empty scaffold (control) or *N*-oleoyl serinol (*N*-OS). One week later, mice were necropsied under ad libitum feeding (Ad Lib.), 12 hours fasted (Fasted), or 12 hours fasted and then re-fed for 3 hours (Re-Fed) to examine meal-related hormone alterations. (**A**) Liver tissue levels of microbe derived *N*-oleoyl serinol (*N*-OS) were quantified by LC-MS/MS analysis. **(B-C)** Hepatic triglycerides and phospholipids lipids, and **(D-F)** plasma lipids (triglycerides, free cholesterol and phospholipids) were determined using commercial enzymatic assay kits. (**G-I**) Plasma levels of other post-prandial incretins and hormones like GLP-1, PYY and glucagon were quantified by immunoassay. **(J-L)** Plasma levels of pro-inflammatory cytokines like IL-6, MCP-1 and TNF-α were quantified by immunoassay. (**M-R**) The relative gene expression for additional metabolism related genes in the liver was quantified by qPCR. (S-X) The relative gene expression for metabolism related genes in the gonadal white adipose tissue (WAT) was quantified by qPCR. Data represent the mean ± S.E.M. from n=5 per group, and statistically significant difference (Student's t-tests) within each feeding status group are denoted by * = p<0.05.

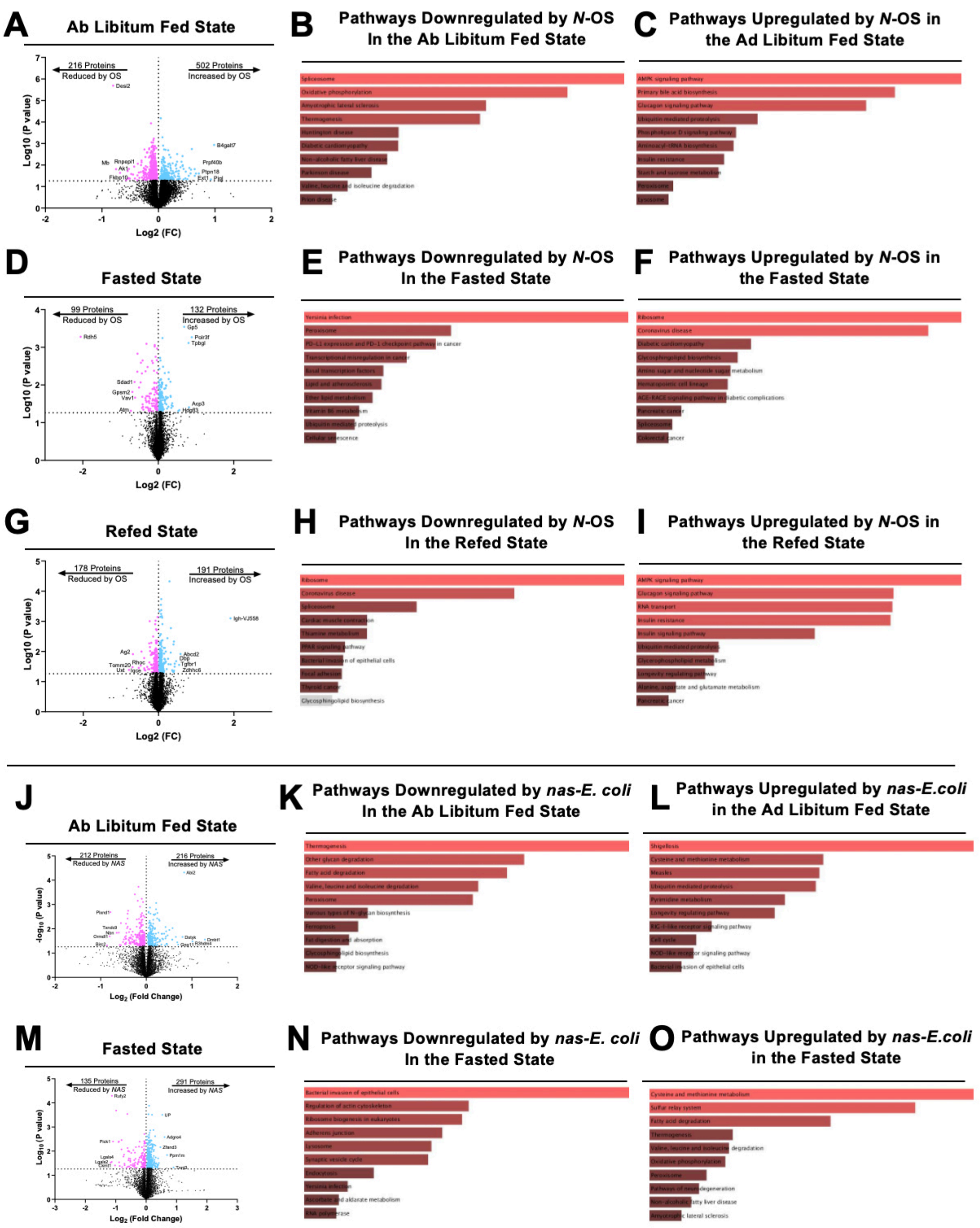


**Figure S2. Global Proteomics Identifies Hepatic Protein Expression Alterations Elicited by Either Exogenous or Gut Microbe-Driven Elevation of *N*-Acyl Amides (*Related to Figure 1*).** (**A-I**) Untargeted proteomics analysis was conducted in liver tissue from animals continuously supplemented with oleoyl serinol via subcutaneous pellets or control (scaffold) control sacrificed in ad libitum (**A-C**), fasted (**D-F**), and refed (**G-I**) states. (**A, D, G**) Volcano plots showing individual proteins downregulated (to the left, significantly reduced proteins are shown in pink) or upregulated (to the right, significantly increased proteins are shown in blue) in animals receiving oleoyl serinol supplementation versus control in ad lib (**A**), fasted (**D**), and refed (**G**) states. (**B, E, H**) Pathway analysis on downregulated proteins was conducted using ErichR and bar charts show the top 10 differentially expressed KEGG pathways for ad lib (**B**), fasted (**E**), and refed (**H**) states. (**C, F, I**) Pathway analysis on upregulated proteins was conducted using ErichR and bar charts show the top 10 differentially expressed KEGG pathways for ad lib (**C**), fasted (**F**), and refed (**I**) states. Significantly altered pathways are shown in shades of red. (J-O) Untargeted proteomics analysis was conducted in liver tissue from germ-free animals recolonized with wild type or *NAS*-expressing *E. coli* in ad libitum-fed, **J-L**) or fasted (**M-O**) states. (**J,M**) Volcano plots showing individual proteins downregulated (to the left, significantly reduced proteins are shown in pink) or upregulated (to the right, significantly increased proteins are shown in blue) in animals recolonized with *NAS*-expressing versus wild type *E. coli* in fed (**J**) or fasted (**M**) states. (**K, N**) Pathway analysis on downregulated proteins was conducted using ErichR and bar charts show the top 10 differentially expressed KEGG pathways for fed (**K**) and fasted (**N**) states. (**L, O**) Pathway analysis on upregulated proteins was conducted using ErichR and bar charts show the top 10 differentially expressed KEGG pathways for fed (**L**) and fasted (**O**) states. Significantly altered pathways are shown in shades of red.

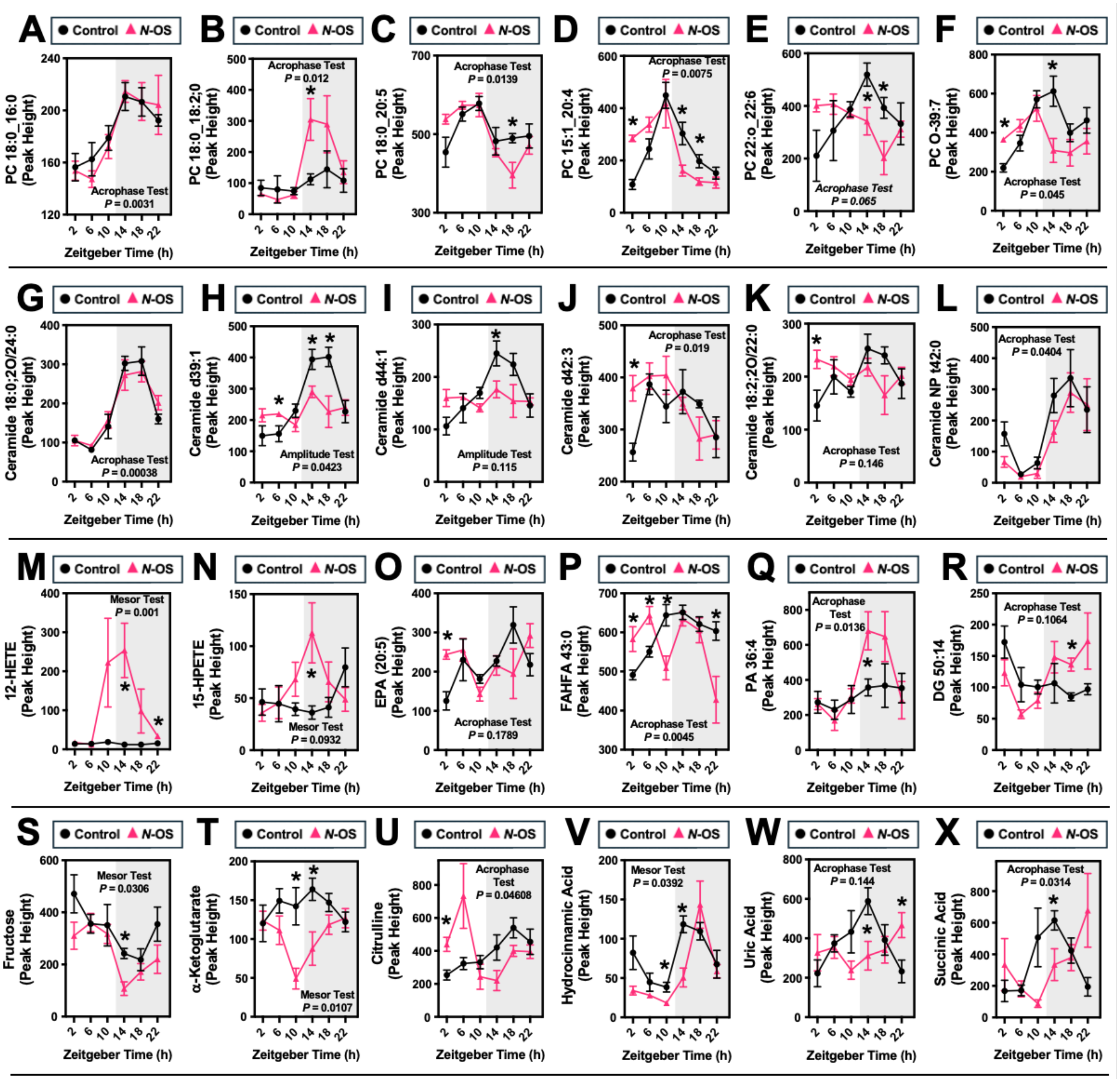


**Figure S3. *N*-Oleoyl Serinol Alters Circadian Rhythms in Systemic Metabolism (*Related to Figure 4*).** Chow-fed C57BL/6J mice were implanted with slow-release pellets containing empty scaffold or *N*-oleoyl serinol (*N*-OS). One week later, mice were necropsied over a 24-hour period (every 4 hours) to examine circadian alterations in metabolic homeostasis. Plasma levels of diverse metabolites were quantified using untargeted lipidomic and metabolomic methods. Data represent the mean -/+ S.E.M. from n=5 per group. Representative group differences were determined using cosinor analyses and *p*-values are provided where there were statistically significant differences between control and *N*-oleoyl serinol treated mice. The entire analysis for circadian cosinor statistics can be found in Table S1. * = statistically significant difference ($p$<0.05) between groups within each ZT time point (Student's t-tests).

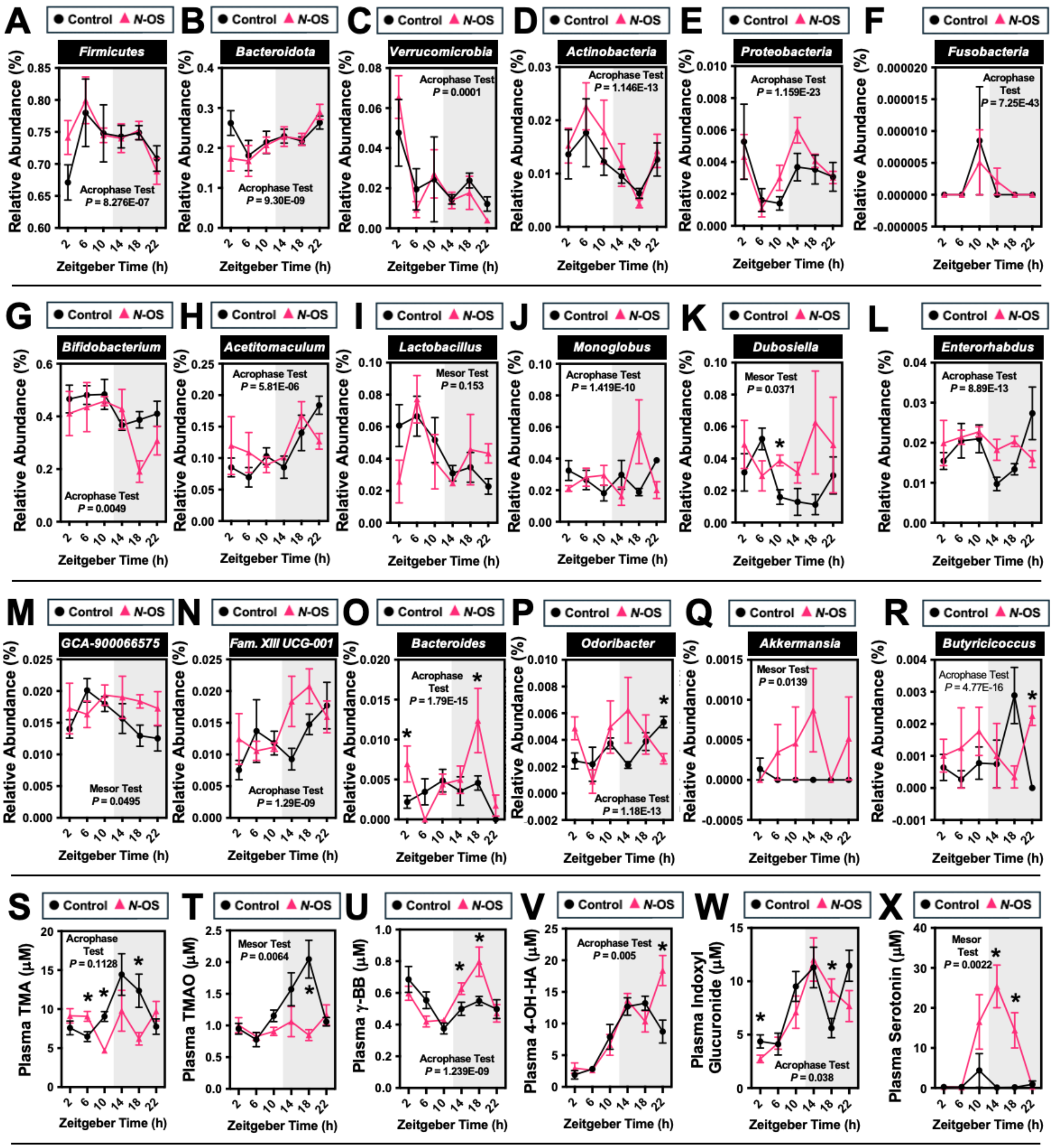


**Figure S4. *N*-Oleoyl Serinol Alters Circadian Rhythms of the Gut Microbiome and Microbe-Derived Metabolites (*Related to Figures 3 & 4*).** Chow-fed C57BL/6J mice were implanted with slow-release pellets containing empty scaffold or *N*-oleoyl serinol (*N*-OS). One week later, mice were necropsied over a 24-hour period (every 4 hours) to examine circadian alterations in metabolic homeostasis. (**A-R**) Cecal microbiome analyses were conducted via sequencing the V4 region of the 16S rRNA (genus level changes are shown). (S-X) Plasma levels of gut microbe-associated metabolites including trimethylamine (TMA), trimethylamine N-oxide (TMAO), γ-butyrobetaine (γBB), 4-hydroxy-hippuic acid (4-OH-HA), indoxyl glucuronide, and serotonin were quantified using stable isotope dilution liquid chromatography tandem mass spectrometry (LC-MS/MS). Data represent the mean -/+ S.E.M. from n=5 per group. Representative group differences were determined using cosinor analyses and *p*-values are provided where there were statistically significant differences between control and *N*-oleoyl serinol treated mice. The entire analysis for circadian cosinor statistics can be found in Table S1. * = statistically significant difference ($p$<0.05) between groups within each ZT time point (Student's t-tests).

**Table S1.** Resources table for methods section – attached as a pdf.

**Table S2.** Cosinor analyses for all circadian studies - attached as a pdf.

**Table S3.** Proteomics analyses – available on request to corresponding and first authors.
