## Supplementary table S1 for "Gut Microbe-Derived *N*-Acyl Serinol Lipids Shape Host Postprandial Metabolic Homeostasis"

**Supplementary Table S1:** Resources table for methods section

| REAGENT or RESOURCE | SOURCE | IDENTIFIER |
|---|---|---|
| Bacterial and virus strains | | |
| *Escherichia coli* MP1 | Gift from Dr. Mark Goulian – University of Pennsylvania | Reference 65 |
| Chemicals, peptides, and recombinant proteins | | |
| *N*-Oleoyl serinol (C18:1 Ser) | Cayman Chemicals | #13637 |
| *N*-Oleoyl serinol-d5 (C18:1 Ser-d5) | C/D/N Isotopes Inc | #D-8112 |
| *N*-Palmitoyl serinol (C16:0 Ser) | Cayman Chemicals | #62175 |
| *N*-Palmitoyl serinol-d5 (C16:0 Ser-d5) | C/D/N Isotopes Inc | #D-8107 |
| *N*-Arachidonoyl serinol (C20:4 Ser) | Cayman Chemicals | #62170 |
| *N*-Oleoyl ethanolamide (C18:1 EA) | Cayman Chemicals | #90265 |
| *N*-Oleoyl ethanolamide-d2 (C18:1 EA-d2) | Cayman Chemicals | #10007823 |
| *N*-Palmitoyl ethanolamide (C16:0 EA) | Cayman Chemicals | #90350 |
| *N*-Palmitoyl ethanolamide-d5 (C16:0 EA-d5) | Cayman Chemicals | #9000573 |
| *N*-Stearoyl ethanolamide (C18:0 EA) | Cayman Chemicals | #90245 |
| *N*-Arachidonoyl ethanolamide (C20:4 EA) | Cayman Chemicals | #90050 |
| *N*-Arachidonoyl ethanolamide-d4 (C20:4 EA-d4) | Cayman Chemicals | #10011178 |
| Isopropanol, Optima™LC/MS Grade | Fisher scientific | A461-4 |
| Hexanes (Optima) | Fisher Scientific | H303-4 |
| Chloroform | Fisher Scientific | C606-4 |
| Acetic Acid, Optima LC-MS | Fisher Scientific | A1131AMP |
| Methanol, Optima LC/MS Grade | Thermo Fisher Scientific | A456-4 |
| LC-MS grade Water | Fisher | 7732-18-5 |
| Critical commercial assays | | |
| Monarch Total RNA Miniprep Kit | New England BioLabs | #T2010S |
| U-Plex Metabolic Combo Mouse Kit | Mesoscale Discovery | #K15297K |
| L-Type Triglyceride M Enzyme Color A (R1) | Fujifilm | #994-02891 |
| L-Type Triglyceride M Enzyme Color B (R2) | Fujifilm | #990-02991 |
| Multi-Lipid Calibrator | Fujifilm | #464-01601 |
| Free Cholesterol E | Fujifilm | #993-02501 |
| Phospholipids C | Fujifilm | #997-01801 |
| Infinity™ Cholesterol Liquid Stable Reagent | Thermo Scientific | #TR13421 |
| Experimental models: Organisms/strains | | |
| Germ/free C57BL/6NTac | Taconic | Stock #: B6-GF-F |
| C57BL/6J | Jackson | #00664 |
| Oligonucleotides | | |
| *Nr1d1* Forward: ATGCCAATCATGCATCAGGT | Sigma | NA |
| *Nr1d1* Reverse: CCCATTGCTGTTAGGTTGGT | Sigma | NA |
| *Cry2* Forward: GCTGGAAGCAGCCGAGGAACC | Sigma | NA |
| *Cry2* Reverse: GGGCTTTGCTCACGGAGCGA | Sigma | NA |
| *Srebf1* Forward: TCTCACTCCCTCTGATGCTAC | Sigma | NA |
| *Srebf1* Reverse: GCAACCACTGGGTCCAATTA | Sigma | NA |
| *Srebf2* Forward: GCGTTCTGGAGACCATGGA | Sigma | NA |
| *Srebf2* Reverse: ACAAAGTTGCTCTGAAAA | Sigma | NA |
| *Bhlhe40* Forward: TGGTGATTTGTCGGGAAGAAA | Sigma | NA |

| | | |
|---|---|---|
| *Bhlhe40* Reverse: ACGGGCACAAGTCTGGAAAC | Sigma | NA |
| *Acly* Forward: CTCACACGGAAGCTCATCAA | Sigma | NA |
| *Acly* Reverse: TCCAGCATTCCACCAGTATTC | Sigma | NA |
| *Acaca* Forward: ACATTCCGAGCAAGGGATAAG | Sigma | NA |
| *Acaca* Reverse: GGGATGGCAGTAAGGTCAAA | Sigma | NA |
| *Hmgcr* Forward: CTTGTGGAATGCCTTGTGATTG | Sigma | NA |
| *Hmgcr* Reverse: AGCCGAAGCAGCACATGAT | Sigma | NA |
| *Cpt1α* Forward: TCCATGCATACCAAAGTGGA | Sigma | NA |
| *Cpt1α* Reverse: TGGTAGGAGAGAGCAGCACCTT | Sigma | NA |
| *Fgf21* Forward: CCTCTAGGTTTCTTTGCCAACAG | Sigma | NA |
| *Fgf21* Reverse: AAGCTGCAGGCCTCAGGAT | Sigma | NA |
| *Fasn* Forward: GTCACCACAGCCTGGACCGC | Sigma | NA |
| *Fasn* Reverse: CTCGCCATAGGTGCCGCCTG | Sigma | NA |
| *Pck1* Forward: GGAAGAACAAGGAGTGGAGAC | Sigma | NA |
| *Pck1* Reverse: CAGGCAGGGTCAATAATGGG | Sigma | NA |
| *G6pd* Forward: AGGTGACCCTAAGCCGGAC | Sigma | NA |
| *G6pd* Reverse: AGGTTTCTTTGGGTAGAAGACCA | Sigma | NA |
| *Acat1* Forward: GGAAGTTGGGTGCCACTTCG | Sigma | NA |
| *Acat1* Reverse: GGTGCTCTCAGATCTTTGG | Sigma | NA |
| *Acat2* Forward: GGGCTGCTGAATTTCACCAT | Sigma | NA |
| *Acat2* Reverse: GAAGAGAAAGGTCCACATCAGGAT | Sigma | NA |
| *Pnpla3* Forward: TCACCTTCGTGTGCAGTCTC | Sigma | NA |
| *Pnpla3* Reverse: CCTGGAGCCCGTCTCTGAT | Sigma | NA |
| *Irs2* Forward: TCCAGGCACTGGAGCTTT | Sigma | NA |
| *Irs2* Reverse: GGCTGGTAGCGCTTCACT | Sigma | NA |
| *Pgc1α* Forward: CCCTGCCATTGTTAAGAC | Sigma | NA |
| *Pgc1α* Reverse: GCTGCTGTTCCTGTTTTC | Sigma | NA |
| *Ucp1* Forward: ACTGCCACACCTCCAGTCATT | Sigma | NA |
| *Ucp1* Reverse: CTTTGCCTCACTCAGGATTGG | Sigma | NA |
| *CycloA* Forward: GCGGCAGGTCCATCTACG | Sigma | NA |
| *CycloA* Reverse: GCCATCCAGCCATTCAGTC | Sigma | NA |
| Software and algorithms | | |
| GraphPad Prism | https://www.graphpad.com/ | Version 10.4.1 |
| JMP Statistical Discovery | https://www.jmp.com/en/home | Version 17.0.0 |
| DAtest package | https://github.com/Russel88/DAtest/wiki/usage#typical-workflow | |
| cosinor | https://cran.r-project.org/web/packages/cosinor/index.html | Version 1.2.3 |
| Other | | |
| qScript | QuantaBio | #95048-100 |
| Fast SYBR Green Master Mix | Applied Biosystems | #4385618 |
