## Supplementary table S2 for "Gut Microbe-Derived *N*-Acyl Serinol Lipids Shape Host Postprandial Metabolic Homeostasis"

**Supplementary Table S2:** Cosinor analyses for all circadian studies

| | | | Group | Zero Amplitude Test | MESOR | Amplitude | Acrophase | Mesor Test | Amplitude Test | Acrophase Test |
|---|---|---|---|---|---|---|---|---|---|---|
| Figure 4: *N*-Oleoyl Serinol Alters Circadian Rhythms in Metabolism | Plasma *N*-acyl serinol (A), metabolic hormones (B-C) and cytokines (D-E) | ***N*-Oleoyl Serinol** | Placebo | **5.84E-04** | 1.06 ± 0.077 | 1.468 ± 0.109 | 0.929 ± 0.265 | **4.11E-08** | 4.28E-01 | **2.81E-02** |
| | | | N-Oleoyl Serinol | 5.57E-01 | 2.75 ± 0.108 | 2.881 ± 0.107 | 1.026 ± 0.818 | | | |
| | | **GLP-1** | Placebo | 1.57E-01 | 0.042 ± 0.013 | 0.074 ± 0.018 | 0.351 ± 0.588 | 9.40E-01 | 7.38E-01 | **5.64E-11** |
| | | | N-Oleoyl Serinol | 5.30E-02 | 0.04 ± 0.018 | 0.091 ± 0.018 | 1.436 ± 0.357 | | | |
| | | **Leptin** | Placebo | **1.63E-05** | 3619.899 ± 203.35 | 5144.349 ± 290.31 | -1.138 ± 0.187 | 6.82E-01 | 8.77E-01 | **2.69E-03** |
| | | | N-Oleoyl Serinol | **8.88E-06** | 3712.333 ± 285.457 | 5460.146 ± 283.319 | -1.432 ± 0.162 | | | |
| | | **TNFa** | Placebo | **1.03E-04** | 4.264 ± 0.225 | 5.827 ± 0.323 | 1.373 ± 0.201 | 4.03E-01 | 9.72E-01 | **3.54E-02** |
| | | | N-Oleoyl Serinol | **9.98E-05** | 4.59 ± 0.316 | 6.189 ± 0.314 | 0.772 ± 0.196 | | | |
| | | **MCP-1** | Placebo | 6.18E-02 | 9.498 ± 2.255 | 12.228 ± 3.144 | -0.161 ± 1.184 | **7.81E-03** | 4.50E-01 | **2.56E-04** |
| | | | N-Oleoyl Serinol | 9.13E-02 | 21.247 ± 3.166 | 30.944 ± 3.142 | 0.539 ± 0.324 | | | |
| | Hepatic Gene Expression (F-J) | **Hepatic *Nr1d1*** | Placebo | **1.86E-04** | 2.102 ± 0.317 | 4.883 ± 0.452 | -1.559 ± 0.16 | 8.28E-02 | 6.02E-01 | **1.38E-03** |
| | | | N-Oleoyl Serinol | **5.56E-06** | 1.295 ± 0.449 | 2.911 ± 0.452 | 1.534 ± 0.275 | | | |
| | | **Hepatic *Cry2*** | Placebo | 1.02E-01 | 3.908 ± 0.452 | 5.736 ± 0.642 | 0.578 ± 0.349 | 1.52E-01 | 7.17E-01 | 2.85E-01 |
| | | | N-Oleoyl Serinol | 9.12E-02 | 2.612 ± 0.635 | 3.5 ± 0.64 | 1.015 ± 0.698 | | | |
| | | **Hepatic *Cpt1a*** | Placebo | 9.11E-02 | 1.5 ± 0.172 | 2.239 ± 0.242 | -0.72 ± 0.329 | **1.43E-02** | 5.61E-01 | 1.32E-01 |
| | | | N-Oleoyl Serinol | 9.94E-02 | 0.908 ± 0.241 | 1.158 ± 0.239 | 1.554 ± 0.956 | | | |
| | | **Hepatic *Srebf2*** | Placebo | 8.11E-02 | 4.533 ± 0.829 | 8.32 ± 1.161 | -0.408 ± 0.313 | **2.43E-02** | 4.89E-01 | 1.63E-01 |
| | | | N-Oleoyl Serinol | 5.96E-01 | 1.711 ± 1.164 | 2.031 ± 1.156 | -0.244 ± 3.614 | | | |
| | | **Hepatic *G6pd*** | Placebo | **9.33E-03** | 3.931 ± 0.523 | 7.189 ± 0.735 | -0.582 ± 0.228 | **2.08E-02** | 2.36E-01 | **3.74E-02** |
| | | | N-Oleoyl Serinol | 1.89E-01 | 1.863 ± 0.733 | 2.489 ± 0.728 | -1.307 ± 1.162 | | | |
| | Plasma phospholipids (K-L), ceramides (M-N) and sugars (O) | **PC 36:2** | Placebo | **4.35E-03** | 1845019548.938 ± 24582609.942 | 1982573315.375 ± 35831544.654 | 1.018 ± 0.248 | 2.35E-01 | 8.12E-01 | **2.64E-02** |
| | | | N-Oleoyl Serinol | **3.50E-04** | 1777379174.448 ± 34155257.969 | 1928080471.868 ± 33477918.157 | -1.356 ± 0.223 | | | |
| | | **PC 34:1** | Placebo | **2.75E-02** | 895003995.963 ± 21235840.696 | 986072111.004 ± 30194296.751 | 0.784 ± 0.332 | 5.48E-01 | 8.30E-01 | 9.36E-02 |
| | | | N-Oleoyl Serinol | **3.42E-03** | 911818326.094 ± 29505232.31 | 1014711601.828 ± 28622758.16 | 0.626 ± 0.285 | | | |
| | | **Cer d36:1** | Placebo | 7.37E-02 | 114777.859 ± 7919.492 | 132346.813 ± 11288.768 | 0.807 ± 0.639 | **2.87E-03** | **3.71E-02** | **1.42E-03** |
| | | | N-Oleoyl Serinol | **5.26E-05** | 149760.398 ± 11003.4 | 221446.046 ± 10661.362 | 0.672 ± 0.153 | | | |

| | | | | | | | | | | |
|---|---|---|---|---|---|---|---|---|---|---|
| | | **Cer d38:1** | Placebo | **9.49E-06** | 326187.856 ± 15976.274 | 463389.005 ± 22943.168 | 0.874 ± 0.164 | 9.98E-01 | 1.63E-01 | **2.19E-02** |
| | | | N-Oleoyl Serinol | 3.36E-01 | 324162.447 ± 22197.552 | 357478.555 ± 21408.082 | 0.955 ± 0.666 | | | |
| | | **Fructose-6-phosphate** | Placebo | 6.87E-01 | 357.39 ± 52.143 | 385.692 ± 75.701 | 0.978 ± 2.564 | **1.54E-05** | 1.47E-01 | **4.23E-02** |
| | | | N-Oleoyl Serinol | **1.66E-03** | 624.279 ± 72.448 | 1000.983 ± 69.975 | 0.825 ± 0.192 | | | |
| Figure S3: *N*-Oleoyl Serinol Alters Circadian Rhythms in Systemic Metabolism | **Phospholipids (A-F)** | **PC 18:0_16:0** | Placebo | **1.23E-04** | 184395.454 ± 4720.167 | 213796.742 ± 6964.625 | 1.159 ± 0.219 | 9.67E-01 | 7.79E-01 | **3.17E-03** |
| | | | N-Oleoyl Serinol | **1.55E-04** | 183167.546 ± 6558.234 | 219432.418 ± 6342.349 | 1.339 ± 0.18 | | | |
| | | **PC 18:0_18:2;O** | Placebo | 2.53E-01 | 99446.389 ± 18046.061 | 132183.159 ± 27060.013 | -1.513 ± 0.739 | 5.22E-02 | 4.03E-01 | **1.22E-02** |
| | | | N-Oleoyl Serinol | **2.55E-04** | 152868.137 ± 25073.329 | 293432.975 ± 24192.382 | 1.191 ± 0.178 | | | |
| | | **PC 18:0_20:5** | Placebo | 8.34E-02 | 507114.923 ± 11045.723 | 549704.564 ± 15593.958 | -0.721 ± 0.371 | 6.95E-01 | 6.80E-02 | **1.39E-02** |
| | | | N-Oleoyl Serinol | **4.03E-07** | 500135.242 ± 15347.008 | 589916.006 ± 14951.644 | -1.532 ± 0.169 | | | |
| | | **PC 35:5\|PC 15:1_20:4** | Placebo | **3.51E-06** | 2451492.104 ± 163388.799 | 3920896.224 ± 220651.933 | -0.257 ± 0.166 | 7.04E-01 | 9.71E-01 | **7.52E-03** |
| | | | N-Oleoyl Serinol | **2.20E-06** | 2369895.926 ± 227013.592 | 3862480.032 ± 223393.045 | -1.245 ± 0.149 | | | |
| | | **PC 44:6\|PC 22:0_22:6** | Placebo | **8.74E-03** | 35780.485 ± 2308.151 | 48828.631 ± 3226.935 | 0.635 ± 0.256 | 9.90E-01 | 6.84E-01 | 6.54E-02 |
| | | | N-Oleoyl Serinol | **4.92E-03** | 33949.957 ± 3206.962 | 42777.378 ± 3121.085 | -1.564 ± 0.36 | | | |
| | | **PC O-39:7** | Placebo | **5.66E-04** | 440781.907 ± 24639.951 | 601021.54 ± 33781.001 | 0.447 ± 0.226 | 1.63E-01 | 5.24E-01 | **4.56E-02** |
| | | | N-Oleoyl Serinol | **4.44E-02** | 378486.722 ± 34234.928 | 463891.3 ± 33692.667 | -1.242 ± 0.393 | | | |
| | **Ceramides (G-L)** | **Cer 18:0;2O/24:0** | Placebo | **4.89E-08** | 181887.998 ± 9601.173 | 305320.38 ± 14147.18 | 1.142 ± 0.106 | 6.25E-01 | 8.05E-01 | **3.80E-04** |
| | | | N-Oleoyl Serinol | **5.34E-08** | 185524.162 ± 13339.94 | 289821.79 ± 12874.925 | 1.216 ± 0.128 | | | |
| | | **Cer d39:1** | Placebo | **3.23E-07** | 257491.839 ± 12494.597 | 400243.285 ± 18263.176 | 1.048 ± 0.121 | 3.73E-01 | **4.23E-02** | **1.23E-03** |
| | | | N-Oleoyl Serinol | 3.57E-01 | 230717.805 ± 17360.084 | 254799.609 ± 16758.875 | 1.235 ± 0.719 | | | |
| | | **Cer d44:1** | Placebo | **2.14E-05** | 1704376.678 ± 75771.976 | 2383004.124 ± 108028.101 | 0.808 ± 0.158 | 6.95E-01 | 1.15E-01 | 7.49E-02 |
| | | | N-Oleoyl Serinol | 9.56E-01 | 1583376.66 ± 105278.138 | 1611961.314 ± 102070.143 | 0.647 ± 3.661 | | | |
| | | **Cer d42:3** | Placebo | **3.04E-02** | 327674.653 ± 12491.762 | 378382.372 ± 16748.08 | 0.083 ± 0.369 | 2.67E-01 | 6.58E-01 | **2.00E-02** |
| | | | N-Oleoyl Serinol | **9.99E-04** | 351357.384 ± 17356.145 | 419819.183 ± 17104.164 | -1.204 ± 0.248 | | | |
| | | **Cer 18:2;2O/22:0** | Placebo | **1.50E-02** | 196368.698 ± 9786.187 | 243889.061 ± 14205.178 | 0.976 ± 0.287 | 3.57E-01 | 4.43E-01 | 1.46E-01 |
| | | | N-Oleoyl Serinol | 2.64E-01 | 205771.59 ± 13597 | 226452.203 ± 13141.351 | 1.307 ± 0.655 | | | |
| | | **Cer-NP t42:0** | Placebo | **6.99E-04** | 183533.517 ± 21041.897 | 335383.47 ± 31536.044 | -1.479 ± 0.186 | 1.63E-01 | 9.59E-01 | **4.05E-02** |

| | | | | | | | | | | |
|---|---|---|---|---|---|---|---|---|---|---|
| | | | N-Oleoyl Serinol | **3.67E-05** | 137120.559 ± 29235.765 | 284440.206 ± 28685.317 | -1.328 ± 0.195 | | | |
| | Other Lipids (M-R) | **12-HETE** | Placebo | 3.17E-01 | 146689.752 ± 163008.254 | 164492.245 ± 243216 | -1.34 ± 12.347 | **1.00E-03** | 1.57E-01 | **8.65E-03** |
| | | | N-Oleoyl Serinol | **1.51E-03** | 1053000.043 ± 226484.86 | 2315434.549 ± 222050.681 | 0.302 ± 0.176 | | | |
| | | **15-HPETE** | Placebo | 1.15E-01 | 48242.522 ± 6467.539 | 63524.463 ± 8771.087 | -0.319 ± 0.628 | 9.32E-02 | 6.81E-01 | 1.53E-01 |
| | | | N-Oleoyl Serinol | **1.23E-02** | 63217.501 ± 8986.045 | 95814.537 ± 8747.613 | 0.509 ± 0.273 | | | |
| | | **FA 20:5** | Placebo | **1.58E-02** | 2098850.132 ± 144050.397 | 2728989.864 ± 214095.748 | 1.266 ± 0.31 | 3.44E-01 | 6.67E-01 | 1.79E-01 |
| | | | N-Oleoyl Serinol | 8.42E-02 | 2255854.448 ± 200144.675 | 2743329.653 ± 198079.784 | 0.028 ± 0.4 | | | |
| | | **FAHFA 43:0** | Placebo | **5.05E-05** | 59409.073 ± 1575.583 | 66726.925 ± 2208.947 | 0.66 ± 0.31 | 2.82E-01 | 9.08E-01 | **4.58E-03** |
| | | | N-Oleoyl Serinol | 2.16E-01 | 57097.809 ± 2189.127 | 62072.291 ± 2186.921 | -0.446 ± 0.424 | | | |
| | | **PA 36:4\|PA 16:0_20:4** | Placebo | 3.20E-01 | 31301.683 ± 3588.702 | 38378.766 ± 5379.743 | 1.493 ± 0.68 | 1.47E-01 | 3.57E-01 | **1.36E-02** |
| | | | N-Oleoyl Serinol | **2.53E-04** | 39529.614 ± 4986.169 | 65449.648 ± 4807.36 | 1.031 ± 0.192 | | | |
| | | **DG 50:14** | Placebo | 6.63E-02 | 1137587.208 ± 99993.683 | 1457283.54 ± 147417.006 | 0.926 ± 0.43 | 7.32E-01 | 6.70E-01 | 1.06E-01 |
| | | | N-Oleoyl Serinol | **6.29E-03** | 1196719.211 ± 136203.954 | 1701126.377 ± 131467.982 | -1.161 ± 0.258 | | | |
| | Sugars/Carbohydrates (S-X) | **Fructose** | Placebo | **1.29E-02** | 33810.344 ± 2119.08 | 45161.145 ± 3063.118 | 0.935 ± 0.261 | **3.06E-02** | 8.94E-01 | **4.94E-03** |
| | | | N-Oleoyl Serinol | **2.07E-04** | 24643.525 ± 2944.265 | 36285.359 ± 2844.324 | 1.282 ± 0.252 | | | |
| | | **α-Ketoglutarate** | Placebo | 1.73E-01 | 13979.008 ± 700.542 | 15892.212 ± 946.519 | 0.264 ± 0.545 | **1.07E-02** | 7.00E-01 | 7.40E-02 |
| | | | N-Oleoyl Serinol | **6.45E-03** | 10369.723 ± 973.339 | 13637.839 ± 970.376 | -0.266 ± 0.288 | | | |
| | | **Citrulline** | Placebo | **6.49E-03** | 3820.407 ± 368.106 | 5108.55 ± 551.627 | 1.473 ± 0.383 | 5.36E-01 | 6.41E-01 | **4.61E-02** |
| | | | N-Oleoyl Serinol | **4.52E-02** | 4102.369 ± 511.449 | 5711.504 ± 493.442 | 0.91 ± 0.317 | | | |
| | | **Hydrocinnamic Acid** | Placebo | **1.75E-02** | 7678.726 ± 729.659 | 10940.343 ± 1094.066 | 1.509 ± 0.3 | **3.92E-02** | 5.55E-01 | **7.40E-03** |
| | | | N-Oleoyl Serinol | **3.40E-04** | 5586.947 ± 1013.793 | 10398.042 ± 990.814 | -1.438 ± 0.208 | | | |
| | | **Uric Acid** | Placebo | **1.81E-03** | 37134.565 ± 2979.435 | 53985.71 ± 4020.283 | 0.243 ± 0.264 | 4.52E-01 | 4.78E-01 | 1.44E-01 |
| | | | N-Oleoyl Serinol | 1.91E-01 | 34074.454 ± 4139.649 | 41627.517 ± 4134.71 | -0.417 ± 0.529 | | | |
| | | **Succinic Acid** | Placebo | **9.26E-04** | 35307.299 ± 4936.097 | 59288.696 ± 6747.467 | 0.415 ± 0.304 | 9.60E-01 | 8.60E-01 | **3.14E-02** |
| | | | N-Oleoyl Serinol | **2.00E-02** | 33326.258 ± 6858.249 | 56920.819 ± 6838.723 | -0.77 ± 0.281 | | | |
| **Figure S4: *N*-Oleoyl** | **16S Phylum Alterations (A-F)** | ***Firmicutes*** | Placebo | 2.18E-01 | 0.732 ± 0.012 | 0.764 ± 0.016 | -0.077 ± 0.518 | 3.53E-01 | 9.74E-01 | **8.28E-07** |
| | | | N-Oleoyl Serinol | 1.26E-01 | 0.744 ± 0.016 | 0.774 ± 0.016 | -1.01 ± 0.541 | | | |
| | | ***Bacteroidota*** | Placebo | 1.80E-01 | 0.229 ± 0.011 | 0.256 ± 0.015 | -0.499 ± 0.569 | 2.74E-01 | 8.40E-01 | **9.30E-09** |

| | | | | | | | | | |
|---|---|---|---|---|---|---|---|---|---|
| | | N-Oleoyl Serinol | **4.32E-02** | 0.215 ± 0.015 | 0.256 ± 0.015 | -1.374 ± 0.36 | | | |
| | *Verrucomicrobiota* | Placebo | 4.54E-01 | 0.024 ± 0.005 | 0.033 ± 0.007 | 0.84 ± 0.739 | 7.35E-01 | 9.32E-01 | **1.40E-04** |
| | | N-Oleoyl Serinol | 1.39E-01 | 0.023 ± 0.007 | 0.036 ± 0.007 | 0.869 ± 0.521 | | | |
| | *Actinobacteria* | Placebo | 8.33E-02 | 0.012 ± 0.001 | 0.016 ± 0.002 | 1.276 ± 0.46 | 2.15E-01 | 5.83E-01 | **1.15E-13** |
| | | N-Oleoyl Serinol | **9.68E-03** | 0.014 ± 0.002 | 0.022 ± 0.002 | 1.567 ± 0.284 | | | |
| | *Proteobacteria* | Placebo | 3.10E-01 | 0.003 ± 0 | 0.004 ± 0.001 | -0.568 ± 0.569 | 4.37E-01 | 8.67E-01 | **1.16E-23** |
| | | N-Oleoyl Serinol | **4.57E-02** | 0.004 ± 0.001 | 0.005 ± 0.001 | 1.214 ± 0.458 | | | |
| | *Fusobacteria* | Placebo | 3.73E-01 | 0 ± 0 | 0 ± 0 | -0.574 ± 0.59 | 8.98E-01 | 9.32E-01 | **7.26E-43** |
| | | N-Oleoyl Serinol | 2.58E-01 | 0 ± 0 | 0 ± 0 | -0.238 ± 0.79 | | | |
| **16S Genus Alterations (G-R)** | *Bifidobacterium* | Placebo | 1.04E-01 | 0.435 ± 0.021 | 0.494 ± 0.029 | 1.521 ± 0.517 | 1.28E-01 | 4.70E-01 | **4.97E-03** |
| | | N-Oleoyl Serinol | **2.03E-02** | 0.374 ± 0.031 | 0.484 ± 0.032 | -1.166 ± 0.303 | | | |
| | *Acetitomaculum* | Placebo | **3.17E-02** | 0.108 ± 0.009 | 0.144 ± 0.012 | -1.283 ± 0.34 | 5.14E-01 | 9.05E-01 | **5.81E-06** |
| | | N-Oleoyl Serinol | 1.41E-01 | 0.117 ± 0.013 | 0.143 ± 0.013 | -1.001 ± 0.539 | | | |
| | *Lactobacillus* | Placebo | **4.33E-02** | 0.047 ± 0.005 | 0.065 ± 0.007 | 1.416 ± 0.414 | 1.53E-01 | 8.31E-01 | 5.14E-01 |
| | | N-Oleoyl Serinol | 5.95E-01 | 0.042 ± 0.008 | 0.052 ± 0.009 | 1.258 ± 0.803 | | | |
| | *Monoglobus* | Placebo | 1.88E-01 | 0.026 ± 0.003 | 0.033 ± 0.005 | 0.15 ± 0.673 | 4.44E-01 | 9.45E-01 | **1.42E-10** |
| | | N-Oleoyl Serinol | 5.79E-01 | 0.028 ± 0.005 | 0.035 ± 0.005 | 1.11 ± 0.67 | | | |
| | *Dubosiella* | Placebo | **1.75E-02** | 0.024 ± 0.006 | 0.04 ± 0.008 | 1.04 ± 0.538 | **3.71E-02** | 3.41E-01 | **1.09E-07** |
| | | N-Oleoyl Serinol | 6.40E-01 | 0.043 ± 0.009 | 0.054 ± 0.009 | -0.947 ± 0.88 | | | |
| | *Enterorhabdus* | Placebo | 3.62E-01 | 0.017 ± 0.001 | 0.02 ± 0.002 | 1.312 ± 0.629 | 2.25E-01 | 9.14E-01 | **8.89E-13** |
| | | N-Oleoyl Serinol | 2.06E-01 | 0.02 ± 0.002 | 0.022 ± 0.002 | -0.829 ± 0.802 | | | |
| | *GCA-900066575* | Placebo | **5.26E-03** | 0.015 ± 0.001 | 0.019 ± 0.001 | -0.921 ± 0.3 | **4.95E-02** | 5.29E-01 | **2.37E-14** |
| | | N-Oleoyl Serinol | 6.65E-01 | 0.018 ± 0.001 | 0.019 ± 0.001 | 0.463 ± 0.91 | | | |
| | *Family XIII UCG-001* | Placebo | 5.71E-01 | 0.012 ± 0.001 | 0.014 ± 0.001 | 1.387 ± 0.909 | 1.23E-01 | 3.38E-01 | **1.30E-09** |
| | | N-Oleoyl Serinol | **1.21E-02** | 0.015 ± 0.002 | 0.02 ± 0.002 | 1.494 ± 0.309 | | | |
| | *Bacteroides* | Placebo | 1.27E-01 | 0.003 ± 0.001 | 0.005 ± 0.001 | 0.053 ± 0.669 | 8.12E-02 | 3.38E-01 | **1.80E-15** |
| | | N-Oleoyl Serinol | 1.82E-01 | 0.005 ± 0.001 | 0.008 ± 0.001 | 1.263 ± 0.354 | | | |
| | *Odoribacter* | Placebo | 3.48E-01 | 0.003 ± 0 | 0.004 ± 0.001 | -1.482 ± 0.939 | 3.29E-01 | 7.73E-01 | **1.18E-13** |
| | | N-Oleoyl Serinol | 2.71E-01 | 0.004 ± 0.001 | 0.005 ± 0.001 | 0.626 ± 0.436 | | | |

| | | | | | | | | | | |
|---|---|---|---|---|---|---|---|---|---|---|
| | | ***Akkermansia*** | Placebo | 4.22E-01 | 0 ± 0 | 0 ± 0 | 0.38 ± 2.568 | **1.39E-02** | 3.36E-01 | **9.10E-20** |
| | | | N-Oleoyl Serinol | 7.47E-01 | 0 ± 0 | 0.001 ± 0 | 0.374 ± 0.873 | | | |
| | | ***Butyricicoccus*** | Placebo | 5.02E-02 | 0.001 ± 0 | 0.002 ± 0 | 1.475 ± 0.403 | 5.94E-01 | 6.72E-01 | **4.78E-16** |
| | | | N-Oleoyl Serinol | 8.12E-01 | 0.001 ± 0 | 0.002 ± 0 | 0.616 ± 1.401 | | | |
| | **Circulating Microbial Metabolites (S-X)** | **TMA** | Placebo | **1.32E-03** | 9.622 ± 0.625 | 13.332 ± 0.886 | 0.884 ± 0.237 | 6.05E-02 | 2.09E-01 | 1.13E-01 |
| | | | N-Oleoyl Serinol | 3.31E-01 | 8.107 ± 0.877 | 9.379 ± 0.87 | 0.009 ± 0.684 | | | |
| | | **TMAO** | Placebo | **9.73E-05** | 1.259 ± 0.068 | 1.812 ± 0.097 | 1.187 ± 0.172 | **6.43E-03** | 2.51E-01 | **1.44E-03** |
| | | | N-Oleoyl Serinol | 5.71E-01 | 0.969 ± 0.095 | 1.054 ± 0.095 | -0.799 ± 1.114 | | | |
| | | **Butyrobetaine** | Placebo | **3.73E-02** | 0.526 ± 0.025 | 0.616 ± 0.035 | 0.121 ± 0.401 | 3.54E-01 | 6.48E-01 | **1.24E-09** |
| | | | N-Oleoyl Serinol | **2.67E-03** | 0.557 ± 0.035 | 0.695 ± 0.035 | -1.541 ± 0.253 | | | |
| | | **4-OH-hippuric acid** | Placebo | **3.14E-08** | 7.872 ± 0.717 | 14 ± 1.023 | 1.083 ± 0.164 | 1.47E-01 | 8.31E-01 | **5.03E-03** |
| | | | N-Oleoyl Serinol | **3.60E-05** | 9.11 ± 1.007 | 15.428 ± 0.999 | -1.524 ± 0.158 | | | |
| | | **Indoxyl-glucuronide** | Placebo | 6.64E-02 | 7.778 ± 0.614 | 10.148 ± 0.872 | 0.916 ± 0.365 | 4.11E-01 | 4.95E-01 | **3.84E-02** |
| | | | N-Oleoyl Serinol | **1.77E-05** | 7.157 ± 0.863 | 11.309 ± 0.856 | 0.919 ± 0.206 | | | |
| | | **Serotonin** | Placebo | 5.23E-01 | 1.06 ± 1.273 | 2.253 ± 1.792 | -0.625 ± 1.516 | **2.20E-03** | 1.97E-01 | **5.24E-04** |
| | | | N-Oleoyl Serinol | **6.73E-06** | 9.526 ± 1.787 | 22.964 ± 1.773 | 0.482 ± 0.132 | | | |
